# Inducing vulnerability to InhA inhibition restores isoniazid susceptibility in drug resistant *Mycobacterium tuberculosis*

**DOI:** 10.1101/2023.02.06.527416

**Authors:** Gregory A. Harrison, Kevin Cho, Erin R. Wang, Souvik Sarkar, Fredrik Almqvist, Gary J. Patti, Christina L. Stallings

## Abstract

Of the approximately 10 million cases of *Mycobacterium tuberculosis* (*Mtb*) infections each year, over 10% are resistant to the frontline antibiotic isoniazid (INH). INH resistance is predominantly caused by mutations that decrease the activity of the bacterial enzyme KatG, which mediates conversion of the pro-drug INH to its active form INH-NAD. We previously discovered an inhibitor of *Mtb* respiration, C10, that enhances the bactericidal activity of INH, prevents the emergence of INH-resistant mutants, and re-sensitizes a collection of INH-resistant mutants to INH through an unknown mechanism. To investigate the mechanism of action of C10, we exploited the toxicity of high concentrations of C10 to select for resistant mutants. We discovered two mutations that confer resistance to the disruption of energy metabolism and allow for growth of *Mtb* in high C10 concentrations, indicating that growth inhibition by C10 is associated with inhibition of respiration. Using these mutants as well as direct inhibitors of the *Mtb* electron transport chain, we provide evidence that inhibition of energy metabolism by C10 is neither sufficient nor necessary to potentiate killing by INH. Instead, we find that C10 acts downstream of INH-NAD synthesis, causing *Mtb* to become particularly sensitive to inhibition of the INH-NAD target, InhA, without changing the concentration of INH-NAD or the activity of InhA, the two predominant mechanisms of potentiating INH. Our studies revealed that there exists a vulnerability in *Mtb* that can be exploited to render *Mtb* sensitive to otherwise subinhibitory concentrations of InhA inhibitor.

**Significance:** Isoniazid (INH) is a critical frontline antibiotic to treat *Mycobacterium tuberculosis* (*Mtb*) infections. INH efficacy is limited by its suboptimal penetration of the *Mtb*-containing lesion and by the prevalence of clinical INH-resistance. We previously discovered a compound, C10, that enhances the bactericidal activity of INH, prevents the emergence of INH-resistant mutants, and re-sensitizes a set of INH-resistant mutants to INH. Resistance is typically mediated by *katG* mutations that decrease the activation of INH, which is required for INH to inhibit the essential enzyme InhA. Our current work demonstrates that C10 re-sensitizes INH-resistant *katG*-hypomorphs without enhancing the activation of INH. We furthermore show that C10 causes *Mtb* to become particularly vulnerable to InhA inhibition without compromising InhA activity on its own. Therefore, C10 represents a novel strategy to curtail the development of INH resistance and to sensitize *Mtb* to sub-lethal doses of INH, such as those achieved at the infection site.

## Introduction

The disease tuberculosis (TB), caused by *Mycobacterium tuberculosis* (*Mtb*), remains a global health threat. As of 2019, TB was reported to be the 13^th^ leading cause of death worldwide, and the current COVID-19 pandemic has exacerbated challenges in TB disease surveillance and global control efforts (1, 2). A major obstacle in the treatment of *Mtb* infections is that the sterilizing activity of antibiotics is slow and sometimes incomplete at the site of infection due to several contributing factors. The penetration of antibiotics into the *Mtb* lesion can be limited, causing *Mtb* to be exposed to fluctuating and often subinhibitory concentrations of antibiotic. In addition, *Mtb* has the propensity to develop phenotypically drug tolerant populations in the host, which allows a population of *Mtb* to persist in spite of exposure to antibiotic (3–6). Long treatment regimens are required to overcome the issues of drug penetration and bacterial drug tolerance to ultimately clear the infection. The standard of care for the treatment of active TB lasts 6 months, including a 2-month intensive phase of regular doses of isoniazid (INH), rifampicin, pyrazinamide, and ethambutol followed by a 4-month continuation phase of INH and rifampicin (7). Recently, the World Health Organization approved the recommendation for a shortened 4-month treatment regimen that can be made available to some patients, which includes a 2-month intensive phase of INH, rifapentine, pyrazinamide, and moxifloxacin, followed by 2 months and 1 week of INH, rifapentine, and moxifloxacin (7). As a component of the intensive and continuation phases of both longer and shorter treatment regimens, INH is a critical frontline antibiotic that is the cornerstone of our current anti-TB regimens.

The utility of INH for the treatment of *Mtb* infections is threatened by the emergence and prevalence of INH resistant mutant strains of *Mtb*. An estimated 10.7% of newly infected and 27.2% of previously treated cases are INH-resistant (8). INH is a prodrug, and resistance is most commonly caused by mutations in the gene *katG*, which encodes the sole bifunctional catalase-peroxidase enzyme in *Mtb* that is also responsible for converting INH to its active form within the bacteria (8–10). KatG is an oxidative defense enzyme, and its typical substrates are H_2_O_2_ and other peroxides (11). However, KatG acts on INH as a non-canonical substrate to generate a radical intermediate of INH (12), which spontaneously reacts with and attaches to the abundant cofactor NAD(H), forming INH-NAD (10, 13). The INH-NAD adduct, the activated form of INH, inhibits the enzyme InhA (10, 13, 14), which is the enoyl-acyl carrier protein reductase enzyme that functions in the fatty acid synthase II (FAS-II) system (15, 16). InhA is required for the FAS-II system to elongate the shorter fatty acids synthesized by FAS-I to generate long lipid precursors that are subsequently converted to mycolic acids (MAs) through a multi-step process. MAs are an essential structural component of the outermost layer of the *Mtb* cell envelope and, therefore, by inhibiting InhA, INH-NAD compromises the integrity of the *Mtb* cell envelope, leading to growth inhibition and death (17, 18).

Identifying ways to enhance the antibacterial activity of INH has the potential to greatly improve the standard of care for TB. To this end, we recently reported the identification of the bicyclic 2-pyridone compound C10 as a potentiator of INH activity in *Mtb* (19). At concentrations that on its own do not inhibit growth, C10 promotes killing by INH and prevents the emergence of spontaneous INH-resistant mutants (19). Whereas high-level resistance to INH mediated by mutations in *katG* is generally considered to render INH ineffective (20), we discovered that C10 was able to re-sensitize multiple INH-resistant *katG* mutants to inhibition by INH, which had previously not been thought to be possible. The ability of C10 to potentiate INH activity in both WT and INH-resistant *katG* mutant strains of *Mtb* demonstrates that there is a vulnerability in the bacteria that can be exploited to enhance the antimicrobial activity of INH and even circumvent INH resistance (19). Understanding the target and mechanism of action of C10 could lead to the discovery of novel therapeutic approaches that can be used in the clinic to disarm INH resistance in *katG* mutants.

In previous work, we performed RNA-sequencing on C10-treated *Mtb* and found that C10 induces a transcriptional signature consistent with inhibition of respiration (19). We subsequently demonstrated that C10 blocked *Mtb* oxygen consumption and decreased bacterial ATP levels, suggesting that C10 disrupts *Mtb* energy metabolism (19). In the current study, we aimed to determine how C10 potentiates killing by INH and whether this potentiation is linked to inhibition of *Mtb* energy metabolism. We use a combination of forward genetics and chemical biology to reveal that the inhibition of respiration by C10 is not required for C10 to potentiate INH, thereby uncoupling these two effects of C10 on *Mtb* physiology. Instead, we present evidence that C10 restores INH susceptibility in a subset of resistant mutants by enhancing the bacterial vulnerability to InhA inhibition. Our findings reveal potential strategies to improve the efficacy of INH as well as other antibiotics that target mycolic acid metabolism in *Mtb*.

## Results

### Isolation of mutants that are resistant to C10

To better understand the mechanism of action of C10 (Fig. 1A), we chose a forward genetic approach and isolated mutants that are resistant to C10, with the goal of identifying mutations in genes linked to the mechanism of action of C10. Previous studies used 25μM of C10 to disrupt *Mtb* energy homeostasis and deplete bacterial ATP (19). But 25μM of C10 only results in a modest decrease in *Mtb* growth (19). Therefore, to select for C10 resistant mutants, we first determined the concentration of C10 that was sufficient to inhibit growth of wild-type (WT) *Mtb*. By increasing the C10 concentration above 25μM, we found that C10 caused dose-dependent inhibition of *Mtb* growth both in liquid media and on agar plates (Fig. 1B-C), consistent with our previous studies (19). 200μM C10 completely inhibited *Mtb* growth in both conditions, so we used this concentration to select for resistant mutants. By spreading *Mtb* on agar containing 200μM C10 and allowing these agar plates to incubate for a total of 9 weeks, we eventually observed the emergence of spontaneous resistant colonies. We isolated 11 resistant mutants and performed whole genome sequencing to identify mutations that confer the C10-resistance. We found that each resistant strain harbored one of 3 different mutations (Fig. 1D). To probe how these mutations impact C10 sensitivity, we chose representative strains that each harbor one of these 3 mutations as the sole nucleotide change that could be identified by whole genome sequencing. Strain GHTB136 harbors an intergenic mutation between the putative S-adenosylmethionine (SAM)-methyl transferase *Rv0731c* and the *secY-adk-mapA* operon, a C to A substitution 69 bp from the start of *Rv0731c* and 92 base pairs (bp) from the start of *secY*. To determine if this mutation impacts the expression level of the neighboring genes we performed qRT-PCR and found that the GHTB136 strain exhibited >100-fold up-regulation of the *Rv0731c* gene and 2-6-fold up-regulation of the *secY-adk-mapA* operon compared to WT (Fig. S1A). The GHTB146 strain harbors an intergenic mutation between the *lpdA*-*glpD2* operon and the uncharacterized gene *Rv3304*, an A to G substitution 12 bp from the start of the *lpdA* coding region. Since *lpdA* is a leaderless transcript (21), this mutation is likely located within the RNA polymerase binding region. The GHTB146 mutant exhibited 4-8-fold up-regulation of the *lpdA*-*glpD2* operon, but no change in expression of the *Rv3304* gene compared to WT (Fig. S1B), suggesting that this mutation enhances *lpdA-glpD2* promoter activity. The third representative strain, GHTB149, harbors a missense mutation within the putative SAM-methyl transferase *Rv0830* that results in the substitution of valine for a leucine residue at position 292 (L292V).

**Figure 1:**
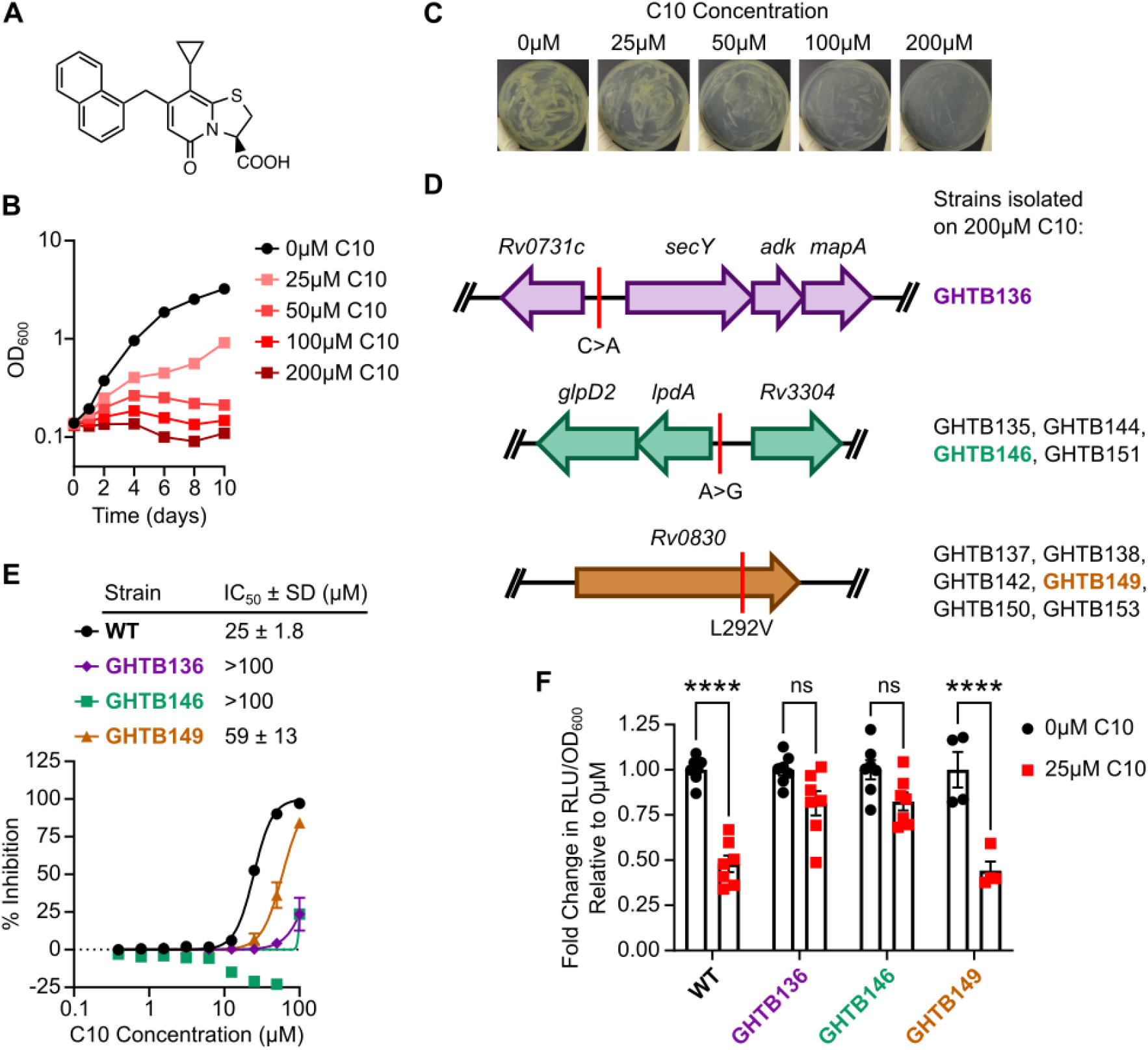
Isolation and characterization of C10-resistant mutants. (A) Chemical structure of C10. (B) WT *Mtb* was cultured in Sauton’s medium containing the indicated concentration of C10, and growth was measured by OD_600_ over time. (C) WT *Mtb* was spread on Sauton’s agar medium containing the indicated concentration of C10 and incubated at 37°C for 3 weeks. (D) Whole genome sequencing of 11 C10-resistant mutants revealed 3 groups of mutants. The mutant loci are depicted along with the resultant nucleotide or amino acid change, and the GHTB strain numbers indicate the mutant isolates that harbored the depicted mutation. Representative strains GHTB136, GHTB146, and GHTB149 were selected for follow up studies. (E) The indicated strain of *Mtb* was cultured in the presence of increasing concentrations of C10 for 1 week, and the % inhibition of Mtb growth and metabolism was determined using the resazurin assay, n=6. (F) The indicated strain of *Mtb* was cultured in Sauton’s liquid medium containing 0 or 25μM C10 for 24 hours before ATP levels were measured by the BacTiter Glo assay. The relative luminescence units (RLU) were normalized to the optical density (OD_600_) of the culture to control for differences in cell density. Fold change in ATP levels were calculated relative to the 0μM C10 control for each strain, n=4-7. A 2-way ANOVA with Tukey’s post test was performed to determine statistically significant differences across samples. Select comparisons are depicted in the figure. ns not significant and **** P<0.0001. For all pairwise comparisons, please see Supplementary Table S1.

To determine how these mutations impact the sensitivity of *Mtb* to C10, we quantified the level of C10 resistance in these strains using a resazurin microplate assay. This assay takes advantage of the redox-sensitive dye resazurin, which is blue in its oxidized form but becomes reduced to the fluorescent pink product resorufin as a result of bacterial growth and metabolism. Therefore, fluorescence can be monitored as a proxy for bacterial growth and metabolism. C10 inhibits WT *Mtb* in this assay with a half-maximal inhibitory concentration (IC_50_) of 25μM (Fig. 1E) (19). The IC_50_ of C10 in GHTB149, which harbors the *Rv0830* L292V mutation, was 59μM (Fig. 1E), a 2.4-fold increase over WT *Mtb*, indicating that this mutant exhibits a low level of resistance to C10. In contrast, the C10 IC_50_ in both GHTB136 and GHTB146 was >100μM (Fig. 1E). Therefore, while this assay confirmed that all 3 representative isolates are resistant to C10, the mutations in GHTB136 and GHTB146 confer a higher level of resistance than the GHTB149 strain.

C10 inhibits respiration and depletes ATP levels in WT *Mtb* (19). Therefore, to directly determine how the mutations in the C10-resistant strains affected ATP-depletion by C10, we cultured WT, GHTB136, GHTB146, and GHTB149 in the presence and absence of 25μM C10 for 24 hours and quantified bacterial ATP levels using a luciferase-based BacTiter Glo assay (Fig. 1F). Similar to WT, the GHTB149 strain exhibited a decrease in bacterial ATP in response to C10, suggesting that the low level of resistance conferred by the *Rv0830* L292V mutation is not sufficient to overcome the depletion of ATP by C10. In contrast, C10 did not significantly decrease the ATP levels in the GHTB136 and GHTB146 strains (Fig. 1F), demonstrating that these mutants are resistant to the ATP-depleting effects of C10. Collectively these findings demonstrate that C10-mediated growth inhibition is linked to ATP depletion, as mutants that maintain ATP levels in the presence of C10 are able to overcome the toxicity of C10. Notably, none of the mutants exhibited an altered level of ATP at baseline (Fig. S2), indicating that these strains do not overcome the effects of C10 simply by harboring increased pools of ATP. Furthermore, the C10-resistant strains were not cross-resistant to compounds known to inhibit ATP synthesis by targeting the electron transport chain (ETC), such as the ATP synthase inhibitor bedaquiline (BDQ) or the protonophore CCCP (Fig. S3). Therefore, GHTB136, GHTB146, and GHTB149 are specifically resistant to the effects of C10, and the mechanism of resistance in these strains does not confer cross-resistance to respiration inhibitors in general.

### C10 potentiates INH independently of its effect on energy homeostasis

Since GHTB136 and GHTB146 overcome the depletion of ATP by C10, we used these mutants to test whether disruption of energy homeostasis by C10 contributes to its ability to enhance the bactericidal activity of INH. We cultured WT *Mtb*, GHTB136, and GHTB146 in media containing 25μM C10 and/or 0.25μg/mL INH and enumerated the colony forming units (CFU) to determine the number of viable bacteria after 10 days of treatment. Similar to our previously reported results, C10 enhanced the bactericidal effect of INH against WT *Mtb*, leading to approximately 2 orders of magnitude fewer CFU/mL after 10 days of treatment compared to INH alone (Fig. 2A) (19). We found that C10 still potentiated the bactericidal activity of INH against GHTB136 and GHTB146 strains to a similar or even greater extent compared to WT (Fig. 2B-C). Therefore, C10 can enhance killing by INH in strains that maintain ATP levels during exposure to C10, demonstrating that ATP depletion is not required for C10 to potentiate INH. In support of this finding, when we examined whether the direct ATP synthase inhibitor BDQ could recapitulate the effect of C10, we found that depletion of bacterial ATP with BDQ did not potentiate killing by INH (Fig. 2D-E). Instead, BDQ in combination with INH resulted in a 1-2-log increase in the number of viable bacteria compared to INH alone. These findings are consistent with previous reports that BDQ and other ETC inhibitors, including Q203 and CCCP, antagonize killing by INH (22–24). Depletion of ATP is, therefore, not sufficient to potentiate INH, indicating that C10 must impart another effect on *Mtb* that is not reversed in the GHTB136 and GHTB146 mutants to elicit the increased sensitivity to INH.

**Figure 2:**
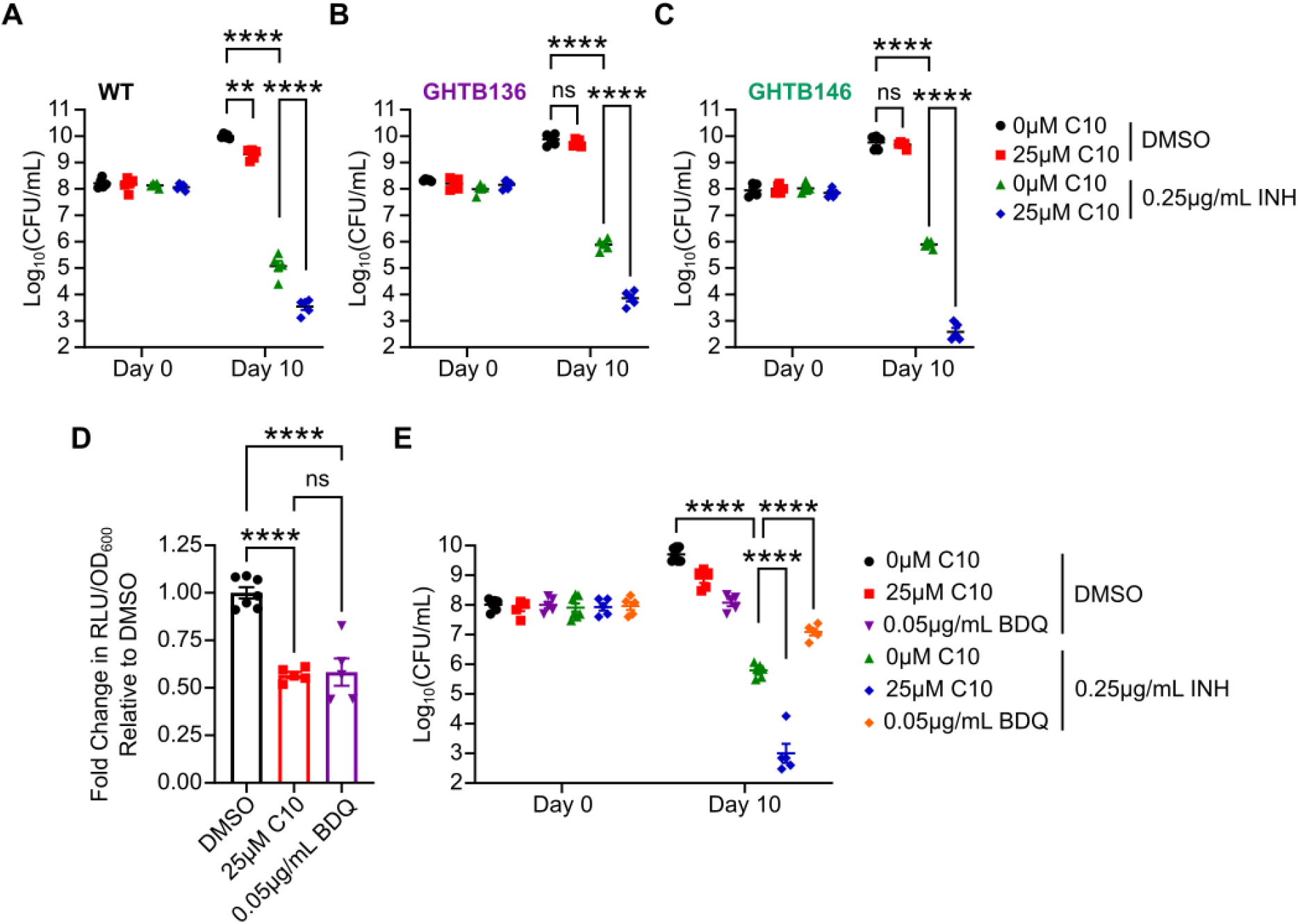
Depletion of ATP by C10 is neither necessary nor sufficient to potentiate the bactericidal activity of INH. (A) WT, (B) GHTB136, or (C) GHTB146 *Mtb* was cultured in Sauton’s liquid medium containing the indicated concentrations of C10 and INH and the CFU/mL were enumerated, n=5. (D) WT *Mtb* was cultured in Sauton’s liquid medium containing 25μM C10 or 0.05μg/mL BDQ for 24 hours before ATP levels were measured by the BacTiter Glo assay. The RLU were normalized to the OD_600_ of the culture to control for differences in cell density. Fold change in ATP levels were calculated relative to the DMSO control, n=5-7. (E) WT *Mtb* was cultured in Sauton’s liquid medium containing the indicated concentrations of C10, BDQ, and/or INH and the CFU/mL were enumerated, n=5-7. A 2-way ANOVA with Tukey’s post test was performed to determine statistically significant differences across samples. Select comparison are depicted in the figure. ns not significant, ** P<0.01, and **** P<0.0001. For all pairwise comparisons, please see Supplementary Table S2.

### Isolation of mutants that are resistant to the combination of C10 and INH

To specifically address how C10 potentiates INH activity in *Mtb*, we sought to identify genes that are required for C10 to potentiate INH by selecting for spontaneous *Mtb* mutants that can grow in the presence of C10 and INH. We had previously been unable to select for mutants that grew in the presence of 25μM C10 and 0.5μg/mL INH (19). Therefore, we decreased the selective pressure by lowering the concentration of INH. We inoculated WT *Mtb* onto agar media containing 25μM C10 and 0.2μg/mL INH, and incubated the bacteria for 4 months at 37°C. We isolated 3 spontaneous mutants that could grow on agar media containing 25μM C10 and 0.2μg/mL INH and performed whole genome sequencing to identify the genetic basis for resistance. Two of the mutant strains harbored large genomic deletions. One strain, GHTB089, was deleted for 27.9 kilobases (kb) of its genome (Δ2145809-2173696), which disrupted 27 annotated genes, including deleting the first 1183 bp of *katG*. The second isolate, GHTB092, was deleted for 38.6 kb of its genome (Δ2132215-2170824), which included the entire *katG* gene and 36 additional annotated genes. In contrast, the third strain harbored a single point mutation when compared to the parental WT strain, a nucleotide change in *katG* that results in an early stop codon at tryptophan 198. We designated this strain *katG*^W198*^.

KatG is a 740-amino acid protein with several residues that are critical for catalase-peroxidase activity, including a heme-coordinating histidine at position 270 and catalytic residues at R104, H108, and W321. The *katG*^W198*^ mutation is predicted to result in truncation of over two thirds of the KatG protein, including several of these essential residues. The selection of the *katG*^W198*^ mutant on media containing INH and C10 was perplexing because in our earlier study we had shown that C10 still potentiates INH in multiple INH-resistant *katG* mutants, including a *katG* mutant that harbors a frameshift mutation at amino acid 6 that results in an early stop codon (*katG*^FS6^) and a *katG* mutant with a single amino acid substitution of a leucine for a tryptophan at position 328 (*katG*^W328L^) (19). To determine if the *katG*^W198*^ mutant was truly unique from the previously studied INH-resistant *katG* mutants, we compared the ability of the *katG*^W198*^, *katG*^FS6^, and *katG*^W328L^ mutants to grow on agar medium containing 0.5μg/mL INH and/or 25μM C10 (Fig. 3A). As expected based on our previous data, the *katG*^FS6^ and *katG*^W328L^ mutants grow well in the presence of either C10 or INH alone, but are re-sensitized to INH in the presence of C10 such that their growth is inhibited on agar containing both C10 and INH together (Fig. 3A). In contrast, the *katG*^W198*^ mutant grew on agar containing C10 alone, INH alone, and the combination (Fig. 3A), demonstrating that this strain is resistant to INH even in the presence of C10.

**Figure 3:**
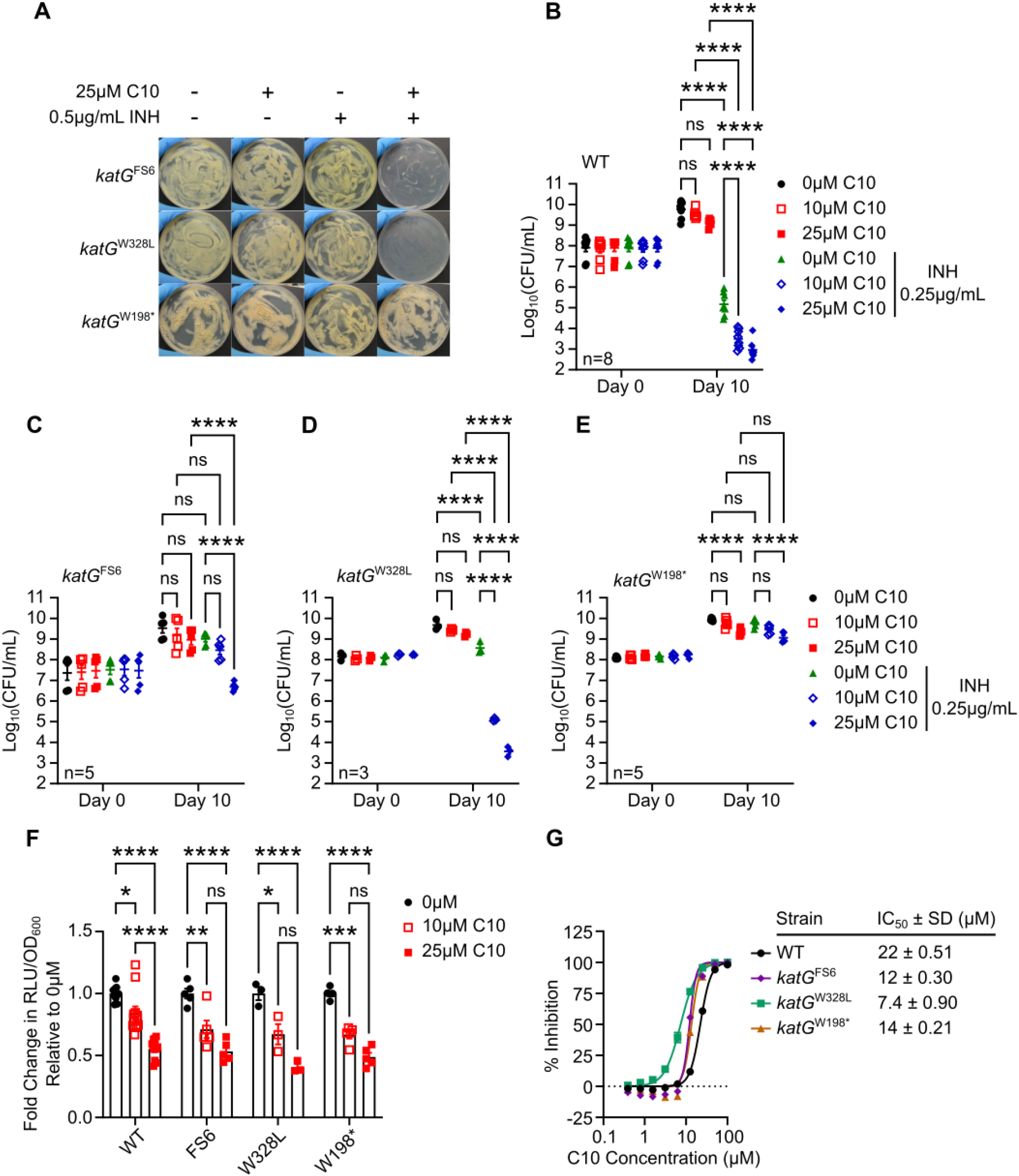
Forward genetic selection results in isolation of a *katG*^W198*^ mutant that is resistant to INH even in the presence of C10. (A) The indicated strain of *Mtb* was spread on Sauton’s agar medium containing 25μM C10 and/or 0.5μg/mL INH and incubated at 37°C for 3 weeks. (B) WT (C) *katG*^FS6^, (D) *katG*^W328L^, and (E) *katG*^W198*^ *Mtb* was cultured in Sauton’s liquid medium containing the indicated concentrations of C10 and INH and the CFU/mL were enumerated. (F) *Mtb* harboring the indicated *katG* allele was cultured in Sauton’s liquid medium containing 0, 10, or 25μM C10 for 24 hours before ATP levels were measured by the BacTiter Glo assay. The RLU were normalized to the OD_600_ of the culture to control for differences in cell density. Fold change in ATP levels were calculated relative to the 0μM C10 control for each strain, n=3-11. (G) The indicated strain of *Mtb* was cultured in the presence of increasing concentrations of C10 for 1 week, and the % inhibition of *Mtb* growth and metabolism was determined using the resazurin assay, n=3. A 2-way ANOVA with Tukey’s post test was performed to determine statistically significant differences across samples. Select comparison are depicted in the figure. ns not significant, * P<0.05, ** P<0.01, *** P<0.001, and **** P<0.0001. For all pairwise comparisons, please see Supplementary Table S3.

We monitored the effects of C10 and INH on the viability of each *katG* mutant strain compared to WT by culturing the bacteria in liquid media with or without C10 and/or INH for 10 days and enumerating CFU (Fig 3B-E). Similar to our previously reported results, while the *katG*^FS6^ mutant can grow in the presence of INH or C10 alone, 25μM C10 in combination with INH caused a significant decrease in the number of viable bacteria compared to C10 or INH alone (Fig. 3C)(19). Similar to the *katG*^FS6^ mutant, the *katG*^W328L^ mutant can grow in the presence of INH or C10 alone, but in combination with INH, 10 or 25μM C10 was able to decrease the number of CFU/mL 3-4 orders of magnitude below the inoculum (Fig. 3D), demonstrating that C10 restores the bactericidal activity of INH against this mutant. In contrast, the CFU/mL of the *katG*^W198*^ mutant increased from day 0 to 10 in cultures treated with C10 alone, INH alone, and the combination (Fig. 3E). 25μM C10 significantly decreased the CFU/mL on day 10 compared to the untreated controls, but this was not enhanced by the addition of INH, indicating that C10 inhibits growth of the *katG*^W198*^ mutant, but is unable to potentiate INH in this strain.

The ability of the *katG*^W198*^ mutant to grow in the presence of C10 and INH could be due to resistance to the C10/INH combination or resistance to C10 specifically. To determine if the *katG*^W198*^ mutant was resistant to C10 activity, we examined the effect of C10 treatment on ATP levels in the *katG*^FS6^, *katG*^W328L^, and *katG*^W198*^ mutants by treating with 10 μM or 25 μM C10 for 24 hours and then measuring ATP levels using the luciferase-based BacTiter Glo assay. C10 treatment caused a similar dose-responsive decrease in ATP in WT *Mtb* and the *katG* mutants as compared to the DMSO-treated cultures (Fig. 3F). C10 also inhibited the *katG* mutants to a similar or even greater extent compared to WT in the resazurin microplate assay, with the IC_50_ of C10 in the WT, *katG*^FS6^, *katG*^W328L^, and *katG*^W198*^ strains being 22μM, 12μM, 7.4μM, and 14μM, respectively (Fig. 3G). These findings demonstrate that C10 is still able to disrupt *Mtb* energy homeostasis in *katG*^W198*^ and, therefore, the loss of INH potentiation in *katG*^W198*^ is not due to insensitivity to C10 activity.

### KatG expression and activity is required for C10 to enhance INH inhibitory activity in *Mtb*

Given the finding that C10 was able to re-sensitize both *katG*^FS6^ and *katG*^W328L^ mutant strains but not *katG*^W198*^ to INH, we hypothesized that there may be a functional difference between the KatG protein variant expressed in the *katG*^W198*^ mutant compared to *katG*^FS6^ and *katG*^W328L^. To begin to investigate this possibility, we probed the expression of KatG protein in each strain by western blot using a monoclonal α-KatG antibody (Fig. 4A). *katG*^W328L^ *Mtb* harbored similar or higher levels of KatG protein compared to WT *Mtb*, whereas *katG*^W198*^ *Mtb* did not express any full length KatG protein, similar to a Δ*katG* strain. In contrast, the *katG*^FS6^ strain expressed a low abundance protein species that was recognized by the α-KatG antibody and migrated slightly faster than the full length KatG protein expressed in WT *Mtb* (Fig. 4A). Upon closer examination of the *katG* mRNA sequence, we noted that there is a putative alternative GUG start codon downstream from the early stop codon introduced by the frameshift mutation in *katG*^FS6^ that may re-initiate translation in the correct frame beginning at codon 23. Therefore, we postulate that the faint band detected by western blot from *katG*^FS6^ cell lysate represents low level expression of a variant KatG protein with a small N-terminal truncation (Δ1-22) that still contains all necessary catalytic residues (Fig. 4A-B).

**Figure 4:**
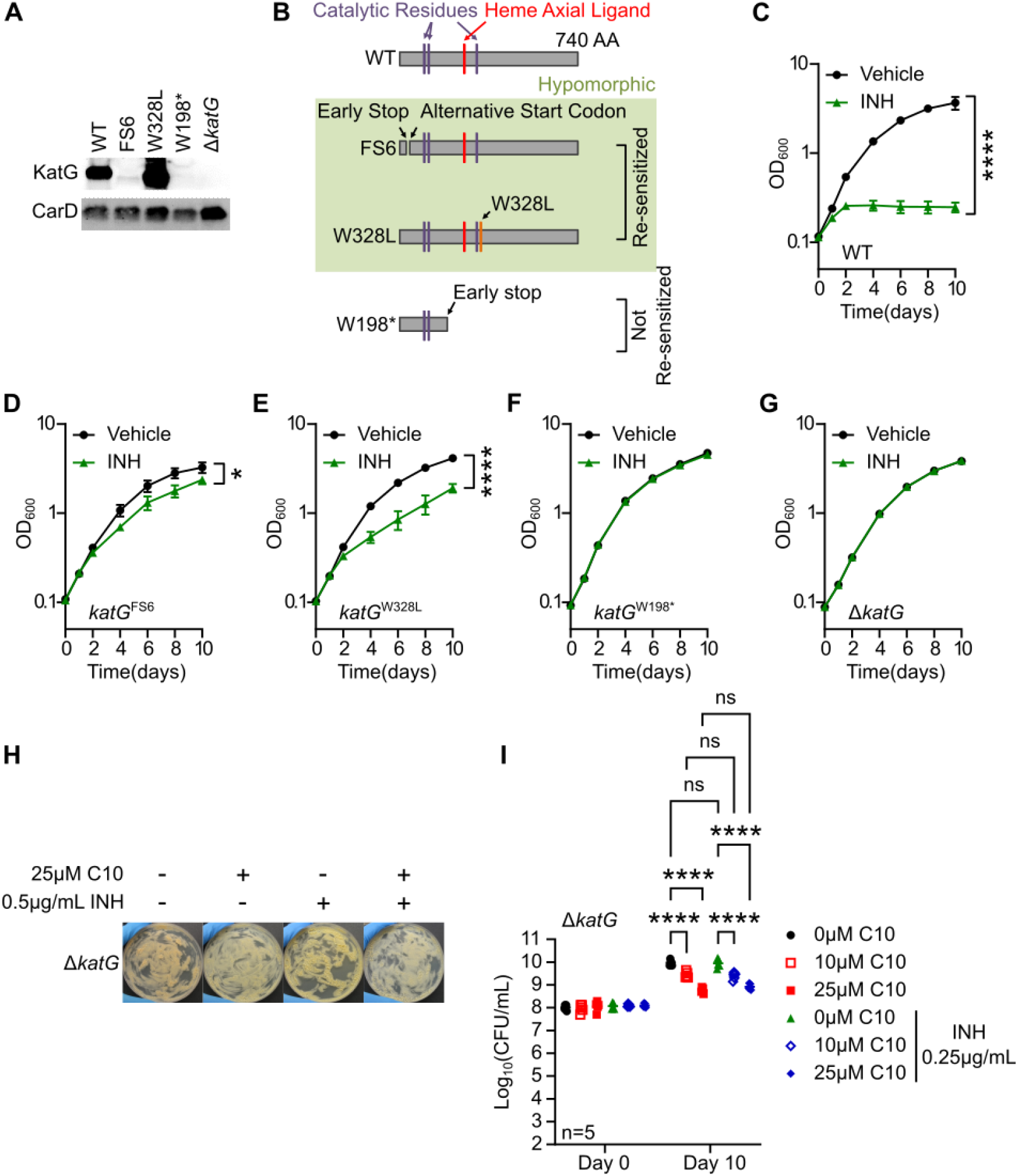
Some level of KatG expression and activity is necessary for C10 to potentiate INH. (A) Western blot against whole cell lysate from the indicated strain of *Mtb*. CarD was used as a loading control in this experiment. (B) Diagram of KatG protein indicating the required catalytic residues in the protein as well as the variant proteins expressed in each *katG* mutant. (C-G) The indicated strain of *Mtb* was treated with or without 0.25μg/mL INH and growth was monitored over time by the optical density (OD_600_), n=4. (H) Δ*katG Mtb* was spread on Sauton agar medium containing 25μM C10 and/or 0.5μg/mL INH and incubated at 37°C for 3 weeks before images were taken. (I) Δ*katG Mtb* was cultured in Sauton’s liquid medium containing 0, 10, or 25μM C10 with or without 0.25μg/mL INH and the CFU/mL were enumerated on 7H11 agar with no antibiotics to determine the number of viable bacteria in each culture at days 0 and 10 of treatment. A 2-way ANOVA with Tukey’s post test was performed to determine statistically significant differences. ns not significant, **** P<0.0001. Relevant comparisons are depicted in the figure, but for all pairwise comparisons please see Supplementary Table S4.

Western blot analysis indicates that the *katG*^FS6^ and *katG*^W328L^ mutants express KatG protein variants that contain all necessary catalytic residues, while *katG*^W198*^ does not (Fig. 4A-B). To determine if the *katG*^FS6^ and *katG*^W328L^ mutants retained KatG catalytic activity, we cultured each strain in liquid media with or without 0.25 μg/mL INH and monitored the growth of the culture by measuring OD_600_ over time. We found that while WT *Mtb* was completely inhibited by 0.25 μg/mL INH (Fig. 4C), the *katG*^FS6^ and *katG*^W328L^ mutants were able to grow in the presence of INH, eventually reaching an OD_600_ over 10-fold above the inoculum (Fig. 4D-E). However, both mutants exhibited a significant decrease in the OD_600_ compared to the untreated controls, indicating that although the *katG*^FS6^ and *katG*^W328L^ mutants exhibit decreased sensitivity to INH, some KatG activity was retained to impart this modest growth inhibition in the presence of INH (Fig. 4D-E). In contrast, the *katG*^W198*^ and Δ*katG* mutants were completely resistant and grew uninhibited in the presence of INH (Fig. 4F-G), supporting that both of these mutations are *katG*-null alleles. Therefore, the *katG*^FS6^ and *katG*^W328L^ mutants that are re-sensitized to INH by C10 are hypomorphic for *katG*, exhibiting decreased KatG activity leading to INH resistance, but retaining a residual level of KatG enzymatic activity (Fig. 4D-E). In contrast, the *katG*^W198*^ mutant that is not re-sensitized to INH by C10 exhibits no KatG activity (Fig. 4F), similar to a *katG*-null strain (Fig. 4G). Based on these data, we hypothesized that some KatG activity is required for C10 to enhance INH sensitivity. To test this hypothesis, we examined whether C10 could sensitize a Δ*katG* mutant to INH. Deletion of *katG* phenocopied the *katG*^W198*^ mutant and enabled *Mtb* to grow on agar and in liquid media containing both C10 and INH (Fig. 4H-I), demonstrating that some KatG expression and activity is required for C10-mediated sensitization to INH.

### C10 induces vulnerability to inhibition by INH without altering KatG activity or INH-NAD levels

Most strategies that renew sensitivity of bacterial pathogens to antibiotics do so by increasing the levels of active antibiotic. Key examples of this are β-lactamase inhibitors that prevent the degradation of β-lactam antibiotics (25) and the recent discovery of SMARt-420, which increases the conversion of ethionamide (ETH) to its active form, ETH-NAD, in *Mtb* (26). In addition, *Mtb* encodes the enzyme CinA that cleaves NAD-drug adducts to promote tolerance to antibiotics like INH and ETH (27), highlighting that regulation of active drug concentration is a major mechanism of modulating drug efficacy. Therefore, we investigated whether C10 sensitizes *Mtb* to INH by promoting KatG activity to enhance the conversion of INH to INH-NAD, thus increasing the levels of INH-NAD in the cell. We monitored the effects of C10 on KatG enzymatic activity *in vitro* by incubating purified KatG protein with H_2_O_2_, a natural substrate of KatG, and monitoring KatG catalase activity as measured by the H_2_O_2_ degradation rate. We found that C10 did not enhance the rate or kinetics of H_2_O_2_ degradation by KatG *in vitro*, indicating that C10 did not directly affect KatG catalase activity (Fig. S4A). We next tested if C10 could specifically promote INH activation by KatG by monitoring the conversion of INH to INH-NAD *in vitro*. Previous work showed that INH activation by KatG occurred most efficiently in the presence of Mn^2+^ (10, 28). Therefore, we incubated KatG protein in buffer containing Mn^2+^ with INH and NAD^+^ in the presence or absence of C10 and monitored the levels of C10, INH, NAD^+^, and INH-NAD by liquid chromatography-mass spectrometry (LC-MS). We found that while the level of C10 remained unchanged over the course of the experiment, INH and NAD^+^ were depleted from the reaction with a concomitant increase in INH-NAD, as expected since INH and NAD^+^ are consumed to produce INH-NAD (Fig. 5A-D). C10 did not impact the rate or level of INH-NAD produced in these conditions. In addition, although we were able to detect the interaction between KatG and INH *in vitro* (Fig. S4B), we were unable to detect a direct interaction between C10 and KatG using a thermal shift assay (Fig. S4C), together supporting that C10 does not directly bind KatG or promote its enzymatic activity *in vitro*.

**Figure 5:**
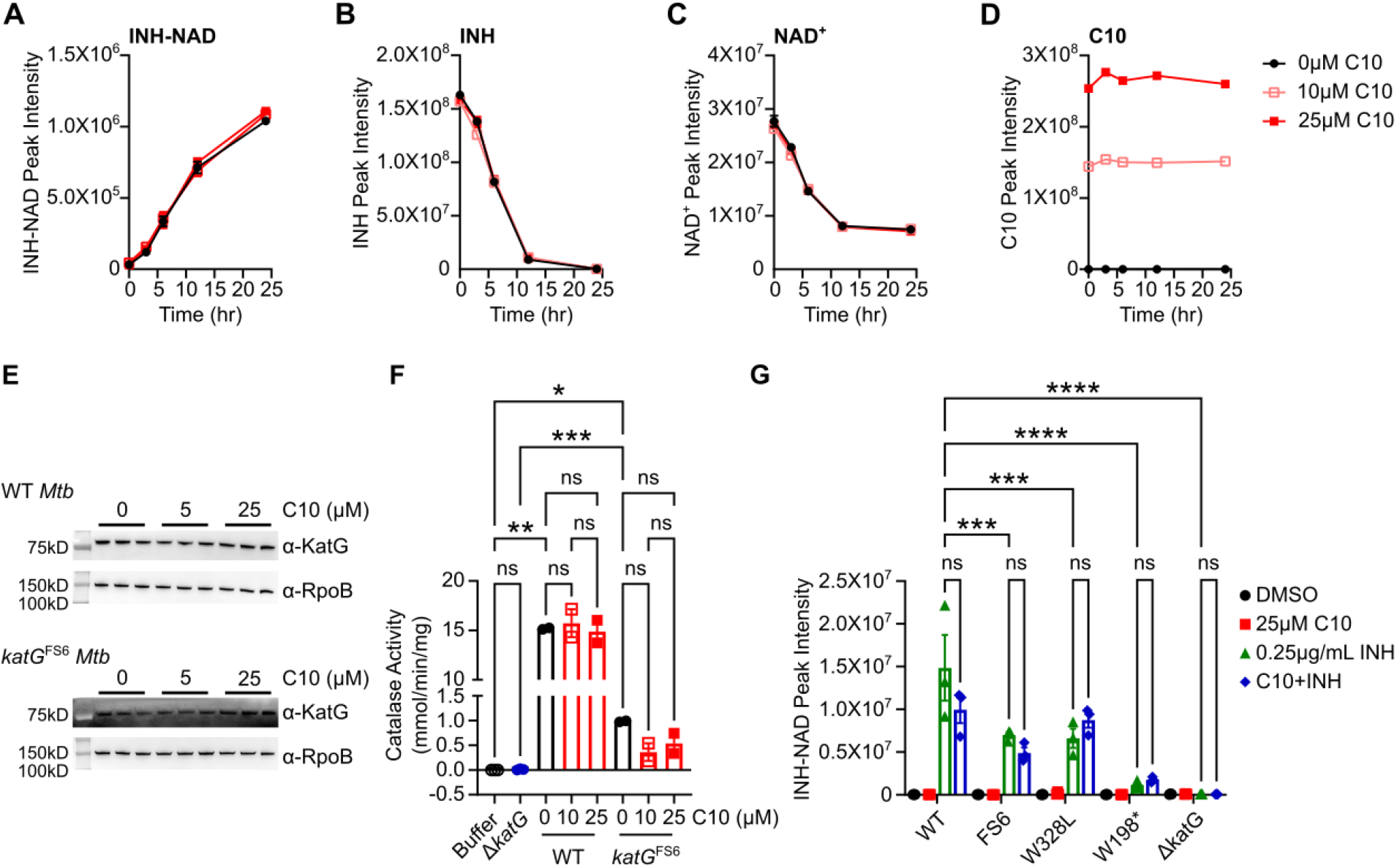
C10 enhances the bacterial sensitivity to INH-NAD without changing its activation. (A-D) Purified KatG protein (200nM) was incubated in 50mM potassium phosphate buffer pH 7.0 containing INH, NAD^+^, and 50μM MnCl_2_ with 0, 10, or 25μM C10, and samples were taken at the indicated time points and analyzed via LC/MS. For each ion, the peak intensity was calculated as the area under the curve for (A) INH-NAD, (B) INH, (C) NAD^+^, and (D) C10, n=3. (E) Either WT (top) or *katG*^FS6^ *Mtb* (bottom) was cultured in Sauton’s liquid medium in the presence of 0, 5, or 25μM C10 for 3 days, and whole cell lysate was collected for a Western blot using a monoclonal α-KatG antibody or an α-RpoB antibody as a loading control, n=3. Note that the KatG blot from the *katG*^FS6^ samples has a high amount of background since the exposure time for this blot is substantially longer than for WT samples due to the very low level of KatG protein in this strain. (F) WT or *katG*^FS6^ *Mtb* was cultured in Sauton’s liquid medium in the presence of 0, 10, or 25μM C10 for 6 days, and whole cell lysate was collected to measure the catalase activity of the sample as a proxy for KatG activity, n=2. Catalase activity is normalized to the amount of total protein in the whole cell lysate sample. Note that lysate from the Δ*katG* mutant, n=3, harbors no detectable amount of catalase activity compared to buffer alone, n=3, demonstrating that the assay is specific for KatG. (G) *Mtb* with the indicated *katG* allele was cultured for 3 days in Sauton’s liquid medium in the presence and absence of 25μM C10 and/or 0.25μg/mL INH, after which polar metabolites were extracted from the culture and analyzed by LC/MS. The area under the curve was calculated by the integration of the peak to determine the peak intensity of INH-NAD in each bacterial sample, n=3. Statistically significant differences were identified by a 2-way ANOVA with Tukey’s post test, and the relevant comparisons are indicated on the graph. ns not significant, *** P<0.001, **** P<0.0001. For all pairwise comparisons see Supplementary Table S5.

To determine if C10 affects KatG activity within the bacterium, we examined whether C10 promotes KatG expression, which could explain how C10 sensitizes WT and *katG* hypomorphic strains to INH but does not sensitize *katG*-null strains. We treated both WT and *katG*^FS6^ *Mtb* with C10 for 3 days and collected whole cell lysate to perform a western blot analysis for KatG. We found that C10 did not increase the protein levels of KatG in these conditions (Fig. 5E). We also monitored the effect of C10 on KatG catalase activity in *Mtb* by treating the bacteria with C10 for 6 days, collecting whole cell lysate, and measuring the H_2_O_2_ degradation rate of the lysate. The Δ*katG* mutant exhibited no H_2_O_2_ degradation in this assay, demonstrating that the assay is specific for KatG (Fig. 5F). While the lysate from *katG*^FS6^ *Mtb* had significantly more activity than the Δ*katG* mutant, this strain exhibited a greater than 10-fold reduction in catalase activity compared to WT, consistent with the *katG*^FS6^ strain being hypomorphic for *katG*. We found that C10 did not change the H_2_O_2_ degradation rate of WT or *katG*^FS6^ *Mtb* (Fig. 5F), confirming that C10 treatment does not enhance the expression or activity level of KatG within *Mtb*. However, this finding did not rule out whether C10 could enhance the levels of INH-NAD within the bacteria without affecting KatG activity. For example, deletion of *cinA*, which encodes an INH-NAD degrading enzyme, leads to elevated INH-NAD accumulation and increased INH sensitivity by decreasing the rate of INH-NAD degradation in *Mtb* (27). To determine if C10 promotes INH-NAD accumulation in *Mtb*, we cultured WT, *katG*^FS6^, and *katG*^W328L^ *Mtb* with and without C10 and/or INH for 3 days, collected aqueous metabolite extracts from the bacteria, and measured the amount of activated INH-NAD in the bacterial extract using LC-MS. We included the *katG*-null mutants *katG*^W198*^ and Δ*katG* as controls. We found that upon treatment with INH alone, WT, *katG*^FS6^, and *katG*^W328L^ strains all accumulated INH-NAD, and the *katG*^FS6^ and *katG*^W328L^ mutants produced significantly decreased levels of INH-NAD compared to WT *Mtb*, confirming that these strains are indeed hypomorphic for *katG* (Fig. 5G). As expected, the *katG*-null mutants *katG*^W198*^ and Δ*katG* were deficient in INH-NAD synthesis (Fig. 5G). C10 did not significantly increase the amount of INH-NAD in any of the strains tested (Fig. 5G) and, therefore, does not affect INH-NAD synthesis or degradation. This data demonstrates that C10 sensitizes *Mtb* to INH through a novel mechanism of action without impacting the levels of INH-NAD in the bacteria.

### C10 sensitizes *Mtb* to direct inhibition of InhA without inhibiting InhA itself

Our data indicates that although the effect of C10 on INH sensitivity relies on KatG activity, C10 does not increase the expression or enzymatic activity of KatG. Based on these findings, we hypothesize that C10 acts downstream of KatG to enhance the antibacterial activity of INH-NAD after it is produced, such that even the *katG* hypomorphic strains that produce lower levels of INH-NAD become inhibited by this lower concentration in the presence of C10. INH-NAD inhibits *Mtb* growth by binding the NADH binding pocket in the enoyl-acyl carrier protein reductase InhA, leading to inhibition of mycolic acid biosynthesis (13, 14, 18, 29, 30). Therefore, it is possible that C10 renders *Mtb* more sensitive to inhibition of InhA. To test this possibility, we cultured WT *Mtb* with C10 and/or NITD-916, a direct inhibitor of InhA that does not require KatG or any other known enzyme for activation (31), and quantified the surviving CFU after 10 days of treatment. Similar to the effect of C10 on INH sensitivity, we found that treating *Mtb* with C10 in combination with NITD-916 caused a significant decrease in the number of surviving bacteria compared to NITD-916 alone, leading to an additional 2-3 orders of magnitude decrease in survival (Fig. 6A). Therefore, C10 increases the bacterial sensitivity to direct InhA inhibition.

**Figure 6:**
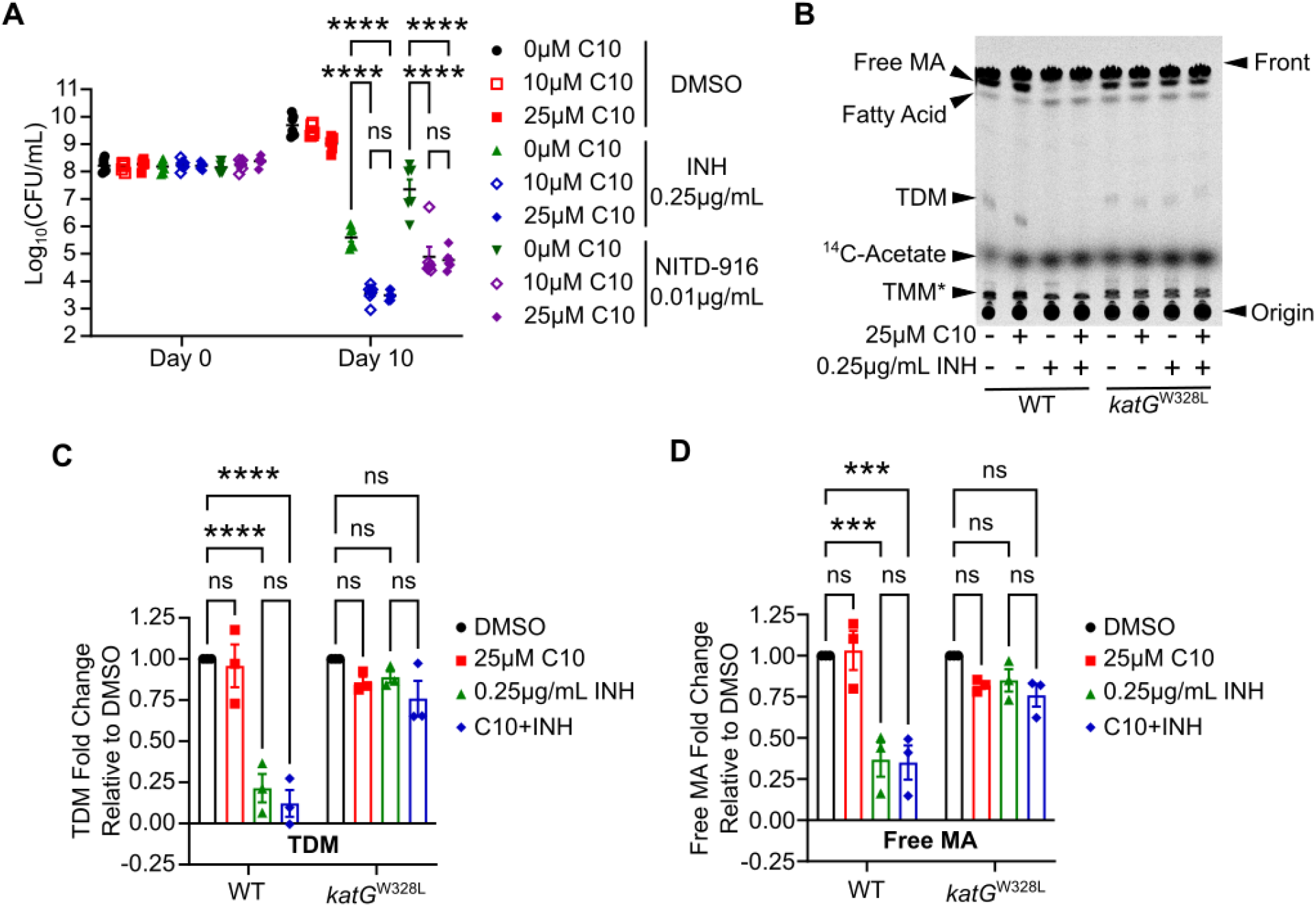
C10 enhances the vulnerability of *Mtb* to InhA inhibition without altering bacterial InhA activity. (A) WT *Mtb* was cultured in Sauton’s liquid medium with the indicated concentrations of C10, INH, and/or NITD-916, and CFU/mL were enumerated on day 0 and 10 of treatment to determine the number of viable bacteria in each sample, n=6. (B) WT or *katG*^W328L^ *Mtb* was cultured in Sauton liquid medium, treated with 25μM C10 and/or 0.25μg/mL INH, and immediately exposed to 2μCi/mL of ^14^C-labeled acetate. After 20 hours, lipids were extracted and analyzed by TLC to measure the *de novo* synthesis of mycolic acids and other lipids. The TLC plate was developed with 75:10:1 Chloroform:Methanol:H_2_O and radioactivity was analyzed by phosphorimaging. Bands corresponding to free mycolic acid (MA), fatty acid, trehalose dimycolate (TDM), and ^14^C-acetate were identified by comigration with a standard. Trehalose monomycolate (TMM) is indicated with a * to emphasize that this lipid is putatively identified, and not correlated with a standard. The plate in (B) is representative of 3 biological replicates, and the standards and additional replicates are shown in Supplementary Figure S5. (C-D) The intensity of bands corresponding to (C) TDM or (D) free MA were quantified in Image J, and normalized to the DMSO sample, with each WT sample being normalized to WT DMSO and each *katG*^W328L^ sample being normalized to *katG*^W328L^ DMSO in order to compare across replicates on separate plates, n=3. Statistically significant differences were determined by 2-way ANOVA and select pairwise comparisons are depicted in the figure. ns not significant, *** P<0.001, **** P<0.0001. For all pairwise comparisons see Supplementary Table S6.

One possible mechanism by which C10 could enhance the susceptibility of *Mtb* to InhA inhibition would be to decrease the expression or activity of InhA, thereby lowering the amount of INH-NAD or NITD-916 required to inhibit this target (32). When we analyzed our previously published RNA-sequencing data from C10-treated cultures, we found that after treatment with 25μM C10 for 48 hours, *inhA* was expressed at 1.1-fold relative to the untreated control (19), demonstrating that C10 does not decrease *inhA* expression at the transcriptional level. We next examined whether C10 compromises InhA activity by culturing *Mtb* with ^14^C-acetate and measuring *de novo* mycolic acid biosynthesis by thin layer chromatography (TLC). In the final steps of mycolic acid biosynthesis, the mycolic acid moiety is coupled to trehalose to form trehalose monomycolate (TMM), which can be transported out of the cell to the envelope (33, 34). The mycolic acid can then be trans-esterified from TMM to the arabinose moieties of arabinogalactan to form the inner leaflet of the mycolic acid layer that is covalently attached to the underlying cell wall layers (35– 37). Alternatively, the mycolic acid can be trans-esterified to a second TMM molecule to form trehalose dimycolate (TDM) (38), which comprises a major component of the freely associated lipid layer that is intercalated within the covalently attached mycolic acids. Free mycolic acid, TMM, and TDM are the primary forms of mycolic acid that are not covalently linked to the cell wall, making them readily extractable and easy to separate by TLC, so we focused our analysis on these mycolic acid species as a read-out for *de novo* biosynthesis. Cultures of WT and *katG*^W328L^ *Mtb* were labeled with ^14^C-acetate for 20 hours in the presence or absence of C10 and/or INH before extracting whole cell lipids and monitoring the incorporation of the ^14^C into mycolic acids by TLC (Fig. 6B-D; Fig. S5). As standards, we used TDM purified from H37Ra (Invivogen), free mycolic acid saponified and extracted from the H37Ra TDM, the representative fatty acid oleic acid, and free ^14^C-acetate. Although we did not have a standard for TMM, we identified a band that we predict corresponds to TMM in our samples since it migrated slower than TDM due to the overall increased polarity and based on the migration pattern reported in published studies (39).

As expected, INH treatment of WT *Mtb* decreased the intensity of bands corresponding to free mycolic acid, TDM, and TMM (Fig. 6B-D; Fig. S5A-D), and significantly increased the intensity of a band that co-migrates with oleic acid (Fig. S5E). We believe this oleic acid co-migrating band represents FAS-I-generated fatty acids that serve as substrates for mycolic acid biosynthesis, thus, it is not surprising that inhibition of InhA would lead to their accumulation (15–17). In contrast, INH treatment of *katG*^W328L^ *Mtb* did not affect the levels of free mycolic acids, TDM, or TMM over the 20 hour period, supporting less efficient InhA inhibition due to the mutation in *katG* and decreased INH-NAD levels (Fig. 6B-D; Fig. S5). Addition of C10 on its own or in combination with INH did not decrease the synthesis of free mycolic acid, TDM, or TMM in the bacteria (Fig. 6B-D; Fig. S5), demonstrating that C10 does not inhibit InhA within the bacteria. While these studies do not rule out that C10 impacts the mycobacterial cell envelope in other ways, our data demonstrate that C10 potentiates the bactericidal effect of InhA inhibition without inhibiting InhA itself or enhancing the ability of INH-NAD to inhibit InhA in *Mtb*.

## Discussion

Together, our findings show that C10 sensitizes *Mtb* to INH without changing the concentration of INH-NAD or decreasing the activity of its target InhA, which are the two predominant mechanisms of potentiating INH reported in the literature thus far (27, 32, 40, 41). Preventing DosR-induced growth arrest and increasing *Mtb* respiration has also been shown to enhance the bactericidal effect of INH (42, 43). However, C10 does not inhibit DosR signaling and decreases *Mtb* respiration (19), indicating that C10 is also not working through these mechanisms. In addition, our studies herein uncouple the effect of C10 on ATP production from the sensitization to INH. Instead, our data supports a model where C10 potentiates INH by making *Mtb* particularly vulnerable to the depletion of cell wall mycolic acids, even in the INH-resistant *katG* hypomorphs that accumulate a significantly lower concentration of INH-NAD. Mycolic acid biosynthesis is essential in mycobacteria and not conserved in non-actinobacteria or eukaryotes, making this process a very attractive target for the development of specific antimycobacterials to treat TB. This is exemplified by INH, which remains a cornerstone of TB treatment, as well as the recent approval of mycolic acid inhibitors delamanid and pretomanid for the treatment of TB and the ongoing development of direct InhA inhibitors, such as NITD-916 (31, 44, 45). In contrast to the potentiation strategies that increase the INH-NAD concentration or decrease InhA expression, which would be specific for INH or InhA inhibitors, respectively, it remains possible that C10 could sensitize *Mtb* more generally to inhibitors of mycolic acid biosynthesis.

The clinical utility of INH is currently being threatened by the increasing rates of INH-resistant TB cases. While resistance to INH can occur through multiple mechanisms, the predominant cause of INH resistance is mutation of *katG*, which accounts for an estimated 78.6% of INH resistant strains (8). Clinically, mutations in *katG* are considered to confer a high level of resistance, often necessitating the use of an alternative treatment regimen. However, the overwhelming majority of these *katG* mutations are not null alleles. The most common resistance variant is an S315T amino acid substitution in KatG that decreases the enzyme’s affinity for INH (8, 46, 47). This S315T mutation and other single amino acid substitutions that are commonly identified in resistant isolates significantly impair the ability of KatG to activate INH, but these mutations do not completely abolish INH-NAD synthesis by the KatG enzyme *in vitro* (47, 48). The complete inactivation of KatG is likely detrimental to *Mtb* survival in the host due to the role of KatG in the oxidative stress response, which could explain why the majority of INH resistant clinical isolates harbor single amino acid substitutions as opposed to more deleterious mutations (49–51). We found that C10 selectively potentiates killing by INH in *katG* mutants that express *katG* and retain its enzymatic activity, revealing that it is possible to rescue the utility of INH against clinically relevant INH-resistant isolates that retain a low level of KatG activity.

INH resistance can also occur upon mutation of the *inhA* locus that either results in increased expression of *inhA* or in expression of an InhA variant that has decreased binding to INH-NAD. An estimated 6.8% of INH-resistant clinical isolates harbor mutations in the *inhA* promoter without a concurrent *katG* mutation (8). Since these mutations confer a low level of INH resistance, patients infected with these strains can be treated with a higher dose of INH to successfully clear the infection (52). Although it has not been directly tested, we anticipate that C10 would potentiate INH efficacy against *inhA* promoter mutants by inducing sensitivity to even low level inhibition of mycolic acid synthesis in these strains.

In addition to resistance, INH efficacy can be limited by subinhibitory concentrations of antibiotic at the site of infection. For instance, a clinical study that quantified the distribution of antibiotic across lung lesions from TB patients within 24 hours of dosing showed that approximately 35% of lesions harbored sub-inhibitory concentrations of INH, likely due to a combination of drug diffusion and host metabolism (5). Therefore, the *Mtb* within these lesions likely experiences fluctuating concentrations of antibiotic that can be sub-inhibitory. By making *Mtb* more sensitive to InhA inhibition, C10 represents a possible strategy to sensitize *Mtb* to even sub-inhibitory concentrations of antibiotic, suggesting that in addition to circumventing INH resistance, C10 could enhance the efficacy of INH at the site of infection, although this remains to be tested. While the precise mechanism by which C10 induces sensitivity to InhA inhibition remains unclear, by deciphering how C10 promotes susceptibility to INH and NITD-916, we could reveal cryptic vulnerabilities in *Mtb* that can be exploited to enhance our current antimicrobial regimen.

## Materials and Methods

### Bacterial strains and growth conditions

*Mtb* Erdman strains (Table 1) were inoculated from a freezer stock into Middlebrook 7H9 liquid medium supplemented with 60 μL/L oleic acid, 5 g/L BSA, 2 g/L dextrose, 0.003 g/L catalase (OADC), 0.5% glycerol, and 0.05% Tween 80 and cultured at 37°C. Actively growing *Mtb* was then inoculated into Sauton’s liquid medium (0.5 g/L KH_2_PO_4_, 0.5 g/L MgSO_4_, 4.0 g/L L-asparagine, 6% glycerol, 0.05 g/L ferric ammonium citrate, 2.0 g/L citric acid, and 0.01% (wt/vol) ZnSO_4_, pH 7.0) supplemented with 0.05% Tween 80 and grown to late-log phase before use in growth curve and survival experiments. The Δ*katG* strain was generated using specialized transduction with the temperature sensitive phage phAE87 engineered to harbor sequence homologous to regions upstream (Erdman nucleotides 2144027-2144776) and downstream (Erdman nucleotides 2146989-2147705) of *katG* and selected on 50μg/mL hygromycin as previously described (53). For *Mtb* growth and survival experiments, *Mtb* was inoculated into roller bottles containing Sauton’s medium supplemented with Tween 80 at a starting OD_600_ of 0.1, and growth was measured by OD_600_ and survival was monitored as CFU/mL. Viable CFU from bacterial cultures were enumerated on Middlebrook 7H11 agar medium supplemented with OADC and 0.5% glycerol and plates were incubated at 37°C with 5% CO_2_ for 2-3 weeks. To select for mutants resistant to both C10 and INH, the equivalent of 0.5mL of OD_600_=1.0 of *Mtb* growing in 7H9+OADC media was spread on Sauton’s agar containing 25μM C10 and 0.2μg/mL INH and incubated at 37°C and 5% CO_2_ for 4 months. Isolated colonies were passaged on agar containing 25μM C10 and 0.2μg/mL INH to ensure that they were resistant before performing whole genome sequencing. For agar growth assays, 0.003 g/L bovine catalase was included in the Sauton’s agar, and the equivalent of 2.5mL of OD_600_=1.0 of *Mtb* was spread on the agar surface to enhance the reproducibility of *Mtb* growth on the agar plates.

**Table 1:**
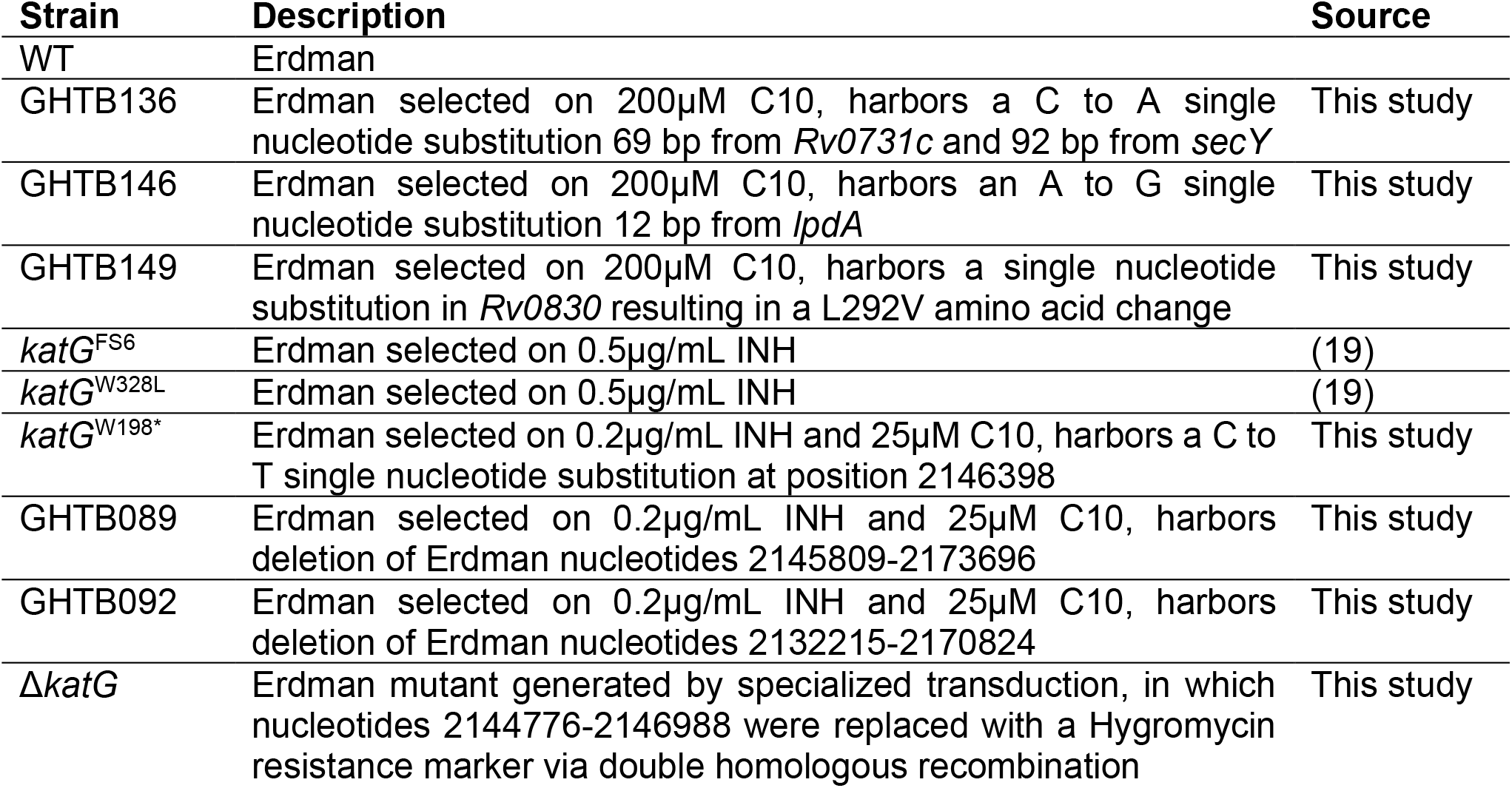
*Mtb* strains used in this study.

### Whole genome sequencing

Genomic DNA was isolated using cetyltrimethylammonium bromide-lysozyme lysis, followed by chloroform-isoamyl alcohol extraction and isopropanol precipitation, as previously described (54). Whole-genome sequencing was performed by use of an Illumina NovaSeq 6000. The identification of single nucleotide polymorphisms was done using SeqMan NGen software (DNASTAR). The genomes were assembled and compared to the genomic DNA from the WT parental control strain, using Integrative Genome Viewer to visualize and confirm changes within regions of interest (55). Whole genome sequencing data is deposited in the Sequence Read Archive (Accession #XXXX).

### Preparation of compounds

C10 was synthesized using previously described methods (56, 57) and prepared as an imidazole salt as described previously (19). Stocks of C10-imidazole were resuspended in DMSO. INH (Sigma) was dissolved in water, and NITD-916 (Sigma) was dissolved in DMSO. In all experiments, the concentration of both DMSO and imidazole were normalized across all samples to ensure that any differences were due to the effect of the indicated compounds and not due to DMSO or imidazole.

### Detection of ATP

*Mtb* growing in Sauton’s media was treated with the indicated concentration of C10 for 24 hours before the OD_600_ of each culture was measured and samples were removed. *Mtb* samples were inactivated at >95°C for 20 min and stored at -20°C until analyzing the ATP levels using the BacTiter Glo assay (Promega) as previously described (19). Samples were diluted 1:10, then mixed 1:1 with the BacTiter Glo reagent in a white, opaque 96-well dish, and the luminescence was read on a Synergy HT plate reader with a 1 second integration. The relative luminescence units (RLU) were calculated by subtracting the luminescence of a media only control from the luminescence value of each sample. The RLU/OD_600_ was determined to account for differences in bacterial density, and the fold-change in each sample was calculated relative to the average of the 0μM C10 control from that experiment to facilitate combining of multiple experiments onto a single graph.

### Resazurin assay

Logarithmically growing *Mtb* was inoculated into Sauton’s medium in 96 well plates with wells containing increasing concentrations of C10. *M. tuberculosis* was inoculated at an ODλ_600_ of 0.0025 in 200 μL per well. The plates were incubated at 37°C in 5% CO_2_ for 1 week, at which point 32.5 μL of a mixture containing an 8:5 ratio of 0.6 mM resazurin (Sigma) dissolved in 1X phosphate-buffered saline to 20% Tween 80 was added, and the production of fluorescent resorufin was measured on a Synergy HT plate reader with excitation λ_ex_ = 530 nm and emission λ_em_ = 590 nm after incubation at 37°C in 5% CO_2_ overnight. For each assay, medium alone served as a negative control, and untreated *Mtb* was included as a positive control. The percent inhibition was calculated as the {[(fluorescence of the positive control − fluorescence of the negative control) − (fluorescence of the sample − fluorescence of the negative control)]/(fluorescence of the positive control − fluorescence of the negative control)} X 100.

### Quantitative reverse transcription PCR (qRT-PCR)

RNA was isolated from WT, GHTB136, and GHTB146 *Mtb* growing in Sauton’s medium using Trizol, and purified by chloroform extraction followed by isopropanol precipitation. cDNA was prepared using the SuperScript III first strand synthesis kit (Invitrogen), and qPCR was performed using a SYBR green kit (Bio-Rad) with gene-specific primers (Table 2) on a CFX96 Real Time System (Bio Rad). The relative expression of genes was calculated using the 2^-ΔΔCt^ method, with *sigA* serving as an internal reference control gene.

**Table 2:**
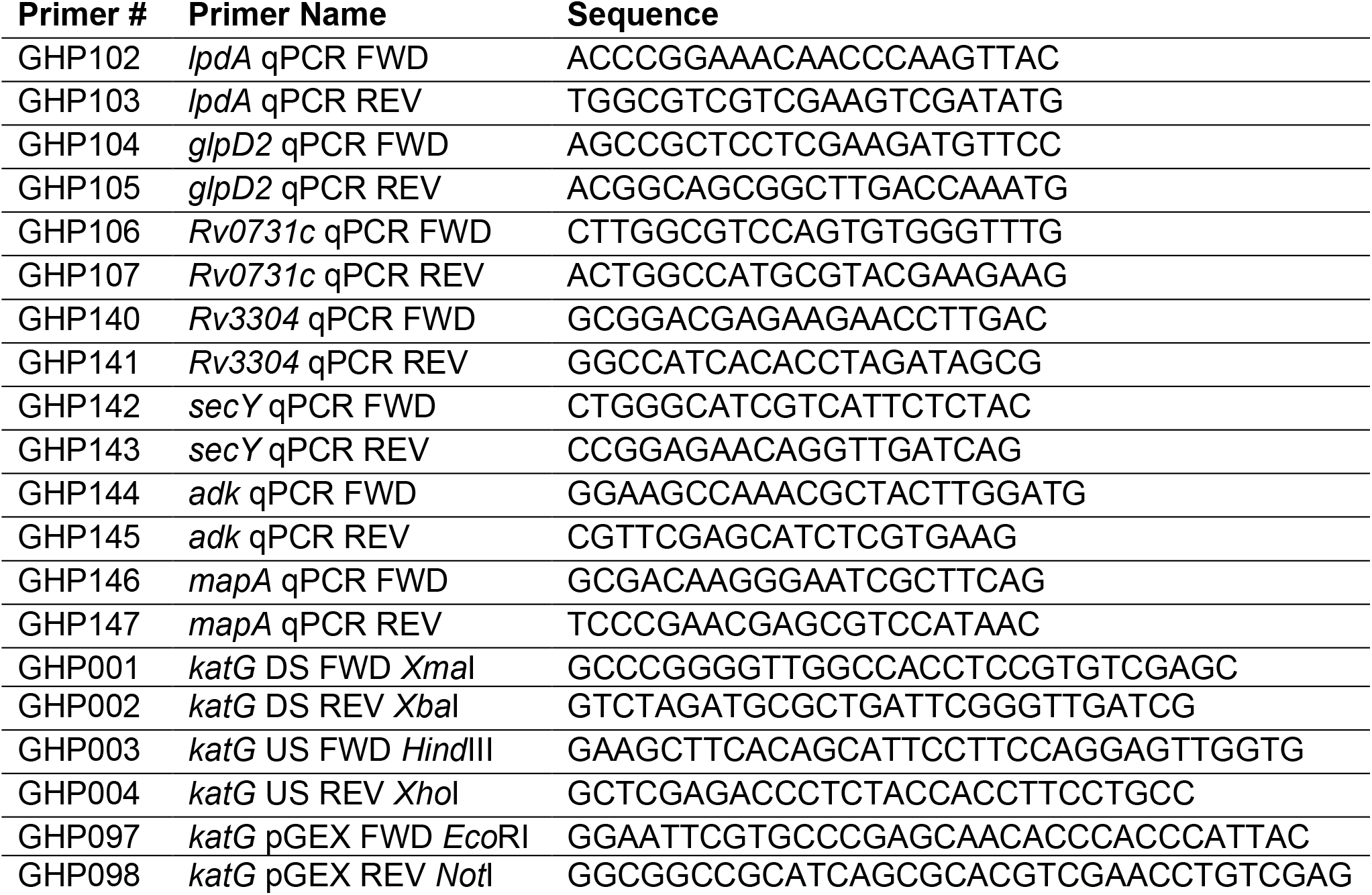
Primers used in this study.

### Western blot for KatG protein

*Mtb* samples were pelleted and resuspended in buffer containing 10mM sodium phosphate pH 8.0, 150mM NaCl, 2mM ETDA, 1mM PMSF, 0.1% NP-40, and a 1X protease inhibitor cocktail (Roche), then lysed by bead beating, and filtered twice through a 0.22μm Spin-X column (Costar) to remove unlysed *Mtb*. SDS-polyacrylamide gel electrophoresis was performed and samples were transferred to a nitrocellulose membrane, after which KatG was detected using a mouse monoclonal α-KatG antibody used at 1:500 dilution (clone IT-57; BEI Resources). Either CarD or RpoB served as a loading control, using a mouse monoclonal α-CarD antibody at 1:2000 dilution (clone 10F05; Memorial Sloan-Kettering Cancer Center) or a mouse monoclonal α-RpoB antibody at 1:1000 dilution (clone 8RB13; Neoclone). The membrane was probed with a goat anti-mouse antibody conjugated to horseradish peroxidase and bands were visualized using the Western Lighting Plus-ECL reagent (Perkin Elmer). When performing the KatG expression analysis in response to C10 treatment, the amount of protein in each sample was measured by BCA (Pierce) and the amount of protein loaded in each lane was normalized to 67ng to facilitate comparisons between samples.

### Purification of *Mtb* KatG protein

The *Mtb katG* coding region was cloned into *Not*I and *Eco*RI sites in the pGEX-6P-1 expression vector to translationally fuse glutathione-S-transferase to the N-terminus of the KatG protein and the expression of the fusion protein was induced in logarithmically growing *E. coli* BL21-DE3 cells by treating 1L of cells with 0.1mM IPTG for 4 hours. Cells were pelleted, resuspended in 20mL 1X PBS containing a 1X protease inhibitor cocktail, and lysed twice in a cell disruptor. Lysate was treated with 9U/mL benzonase (Sigma) and clarified by centrifugation. GST-KatG was purified from the supernatant by incubating lysate overnight with glutathione agarose resin (Goldbio), washed with 300mL 1X PBS, and eluted from the resin by cleaving the KatG protein from the GST tag using PreScission protease in buffer containing 50mM Tris-HCl pH 7.0, 150mM NaCl, 1mM EDTA, and 1mM DTT.

### Thermal shift assay on KatG protein

The melting temperature (T_m_) of KatG in the presence of C10 or INH was determined by differential scanning fluorimetry. Purified *Mtb* KatG protein (1.1μM) was incubated with INH or C10 in 50mM potassium phosphate buffer pH 7.0 containing 50μM MnCl_2_ before mixing samples with SYPRO™ Orange (ThermoFisher) at a final concentration of 1X in a 96-well PCR plate. The plate was incubated for 5 seconds at increasing temperatures in 0.5°C increments from 10-95°C and fluorescence was monitored over time in the HEX channel on a CFX96 Real Time System (Bio Rad). The T_m_ was calculated by fitting curves with a Boltzmann sigmoidal equation in GraphPad Prism, and the ΔT_m_ was calculated as the difference between the T_m_ of each sample and the average of the untreated control samples

### Hydrogen peroxide degradation assay

The catalase activity of either purified KatG or *Mtb* cell lysates was measured in a UV/Vis spectrophotometer using quartz cuvettes. For *in vitro* assays of purified KatG activity, 1.9mL of 50mM potassium phosphate buffer pH 7.0 containing 25nM purified KatG was incubated with or without C10 and/or INH at 30°C for 5min and the sample was used to blank the spectrophotometer before the indicated concentration of H_2_O_2_ was added to initiate the reaction, bringing the final volume of the reaction to 2mL. For samples containing *Mtb* lysate, the samples were bead beat as described above and the protein concentration in the lysate was measured by BCA (Pierce). Samples were normalized such that 25μg of total cell protein was present in each assay sample in 1.9mL of 50mM potassium phosphate buffer pH 7.0, and the reactions were initiated with the addition of H_2_O_2_ to a concentration of 5mM in 2mL final volume. The absorbance at 240nm was read every 10 seconds for 2 min, and the negative slope of the curve was used to calculate the rate of H_2_O_2_ degradation. The absorbance was converted to molarity using the extinction coefficient for H_2_O_2_, ε_240_=43.6M^-1^cm^-1^.

### INH-activation assay and detection of INH-NAD in *Mtb*

The activation of INH was monitored *in vitro* using 200nM purified KatG protein in 50mM potassium phosphate buffer pH 7.0, 50μM NAD^+^, 50μM INH, 50μM MnCl_2_, and the indicated concentration of C10 in a final volume of 1mL. At the indicated time points, 100μL were removed from the reaction and inactivated in 100μL ice cold methanol and stored at -20°C before liquid chromatography/mass spectrometry (LC/MS). To monitor INH activation in live *Mtb*, 50mL cultures of *Mtb* in Sauton’s media without Tween 80 were treated with the indicated concentration of C10 and/or INH for 3 days. To extract polar metabolites, the cultures were pelleted, washed twice in H_2_O, and resuspended in 1.5mL of 2:1 chloroform:methanol in glass conicals. Samples were kept on ice and vortexed each for 1 minute in 20 second intervals, and stored at 4°C overnight. 375μL of H_2_O was added, the samples were vortexed for 1 minute in 20 second intervals, keeping the samples on ice. Samples were incubated at room temperature for 1 hour with constant agitation, and then centrifuged for 10 minutes at 1000 RPM. The top aqueous layer was transferred to a fresh 1.5mL microcentrifuge tube, stored overnight at -20°C, centrifuged for 5 minutes to pellet any insoluble material, and the supernatant was transferred to a fresh tube. INH-NAD was detected using methods similar to those previously described (27, 41, 58) with some modifications. UPLC/MS was performed with an Agilent 1290 Infinity UHPLC system interfaced with an Agilent 6530 QTOF mass spectrometer. Hydrophilic interaction liquid chromatography (HILIC) analysis was performed by using a HILICON iHILIC-(P) Classic column with the following specifications: 100 mm x 2.1 mm, 5 μm. Mobile-phase solvents were composed of A = 20 mM ammonium bicarbonate, 0.1% ammonium hydroxide (adjusted to pH 9.2) and 2.5 μM medronic acid in water:acetonitrile (95:5) and B = 2.5 μM medronic acid in acetonitrile:water (95:5). The column compartment was maintained at 45°C for all experiments. The following linear gradient was applied at a flow rate of 250 μL/min: 0-1 min: 90% B, 1-12 min: 90-35% B, 12-12.5 min: 35-20% B, 12.5-14.5 min: 20% B. The column was re-equilibrated with 20 column volumes of 90% B. The injection volume was 2 μL for all experiments. Data were acquired in both positive and negative ion modes. The mass/charge (*m/z*) and retention times (RT) of the compounds were as follows: INH *m/z*=138.066188, RT=1.87 min; C10 *m/z*=378.11584, 0.92 min; NAD^+^ *m/z*=664.116399, RT=6.12 min; INH-NAD *m/z*=769.137863, RT=5.70 min.

### Measurement of *de novo* lipid synthesis by ^14^C labeling and TLC

To monitor *de novo* mycolic acid biosynthesis, *Mtb* growing in Sauton’s liquid medium was adjusted to OD_600_ 0.5, treated with the indicated concentrations of C10 and/or INH in 1mL final volume, and immediately exposed to 2μCi/mL ^14^C-acetate (Perkin Elmer). After incubation at 37°C for 20 hours, the cells were pelleted, and resuspended in 2:1 chloroform:methanol, vortexed, then samples were pelleted to remove insoluble cell debris, and the supernatant was transferred to a fresh vial. To separate lipid species by TLC, 40μL of the sample was added dropwise to a HPTLC plate coated with silica gel 60 matrix (Sigma). TLC plates were developed in 75:10:1 chloroform:methanol:H_2_O and imaged by phosphorimaging on a Typhoon laser-scanner (Cytiva). The intensity of each band was quantified in ImageJ, normalized to the total intensity in the whole lane, and the fold change was quantified relative to the untreated control for each replicate. To assign putative identities to relevant bands, standards for ^14^C-acetate, TDM, free mycolic acid, and oleic acid were run on each plate. TDM purified from H37Ra (Invivogen) was derivatized to generate a free mycolic acid standard using methods similar to those previously described (59). Briefly, 100μL of 0.5mg/mL TDM in isopropanol was subjected to an alkaline ester hydrolysis by mixing with 2μL H_2_O and 5μL of 10M KOH, and heating to 90°C for 1 hour to ensure efficient saponification. The resulting mycolic acids were purified from the trehalose by neutralizing the reaction with 50μL 1.2M HCl, adding 100μL chloroform, 100 μL H_2_O, vortexing, and separating the organic phase from the aqueous layer to obtain free mycolic acids in chloroform.

## Acknowledgements

We thank Drs. Michael Glickman and Allison Fay for helpful guidance with the ^14^C labeling experiments. We thank the Genome Technology Access Center in the Department of Genetics at Washington University School of Medicine for help with genomic analysis. The Center is partially supported by NCI Cancer Center Support Grant #P30 CA91842 to the Siteman Cancer Center and by ICTS/CTSA Grant# UL1TR002345 from the National Center for Research Resources (NCRR), a component of the National Institutes of Health (NIH), and NIH Roadmap for Medical Research. This publication is solely the responsibility of the authors and does not necessarily represent the official view of NCRR or NIH. This work was supported by a National Science Foundation Graduate Research Fellowship DGE-1745038 (G.A.H.), NIH T32AI007172 (E.R.W.), a Beckman Young Investigator Award from the Arnold and Mabel Beckman Foundation (C.L.S.), an Interdisciplinary Research Initiative grant from the Children’s Discovery Institute of Washington University and St. Louis Children’s Hospital (C.L.S.), NIH R01 AI134847 (C.L.S. and F.A), and NIH R35 ES028365 (G.P.J.). Parts of this project were also supported by the Swedish Research Council 2018-04589 and 2021-05040J (F.A.), the Kempe Foundation SMK-1755 (F.A.), the Erling-Persson Foundation (F.A.), and support under the framework of the JPIAMR – Joint Programming Initiative on Anti-microbial Resistance 2018-00969 (F.A.).

## Author Contributions

G.A.H. and C.L.S. designed the experiments and wrote the manuscript. G.A.H. and E.R.W. performed the experiments with *Mtb*. K.C. performed and analyzed the mass spectrometry experiments with guidance from G.J.P.. S.S. synthesized C10 with guidance from F.A.. All authors contributed to interpreting the data and editing the manuscript.

## Figures and Tables

**Figure S1:**
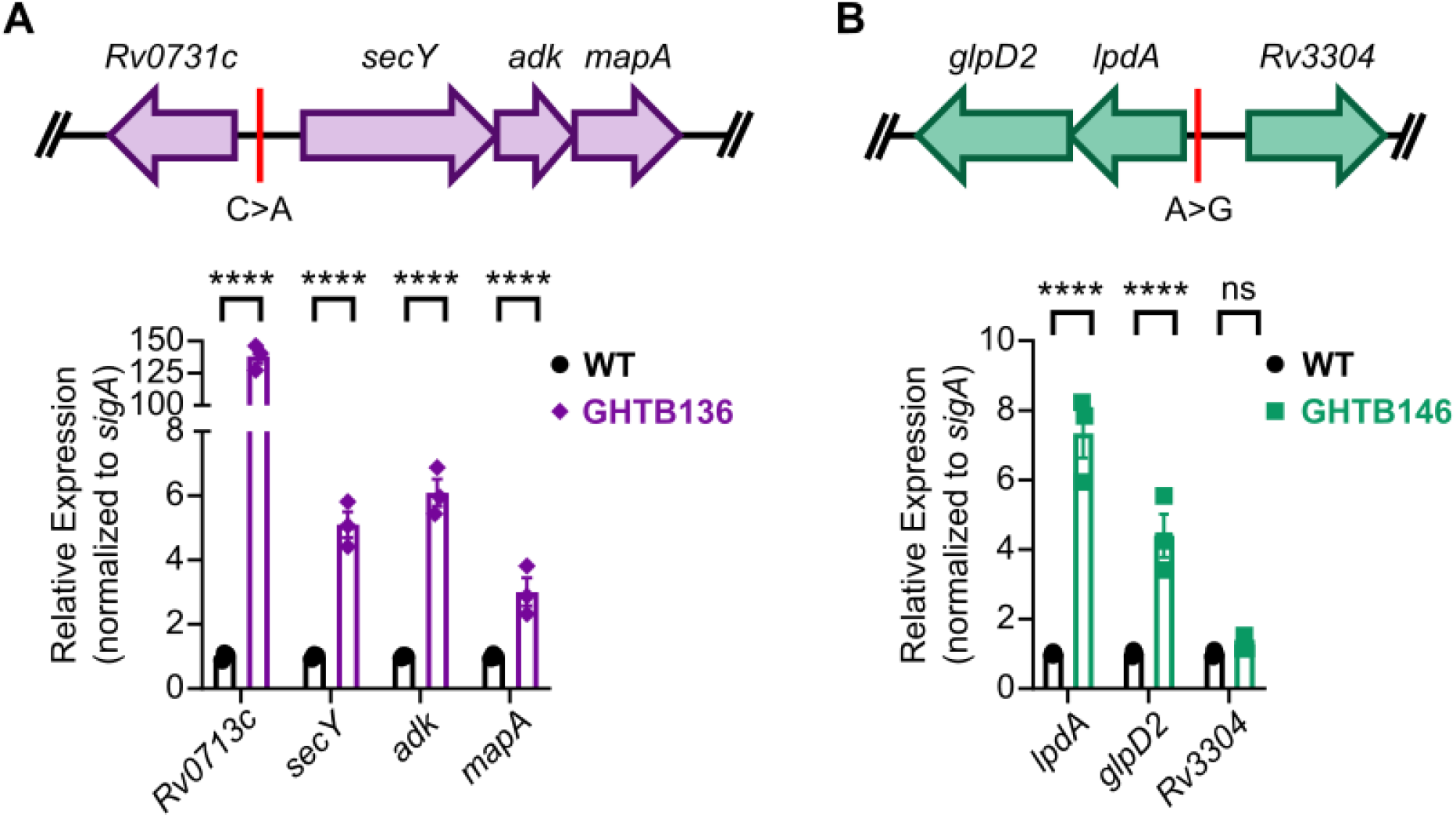
The mutations in GHTB136 and GHTB146 impact the expression of neighboring genes. (A-B) The expression of the indicated genes was monitored in (A) GHTB136 or (B) GHTB146 compared to WT *Mtb* by qRT-PCR, n=3. A map of the gene locus as well as the mutation present in the respective mutant strain is depicted above each graph. The data was log-transformed before performing a 2-way ANOVA with Tukey’s post test to identify statistically significant differences between samples, ****P<0.0001, ns not significant. For all pairwise comparisons see Supplementary Table S7.

**Figure S2:**
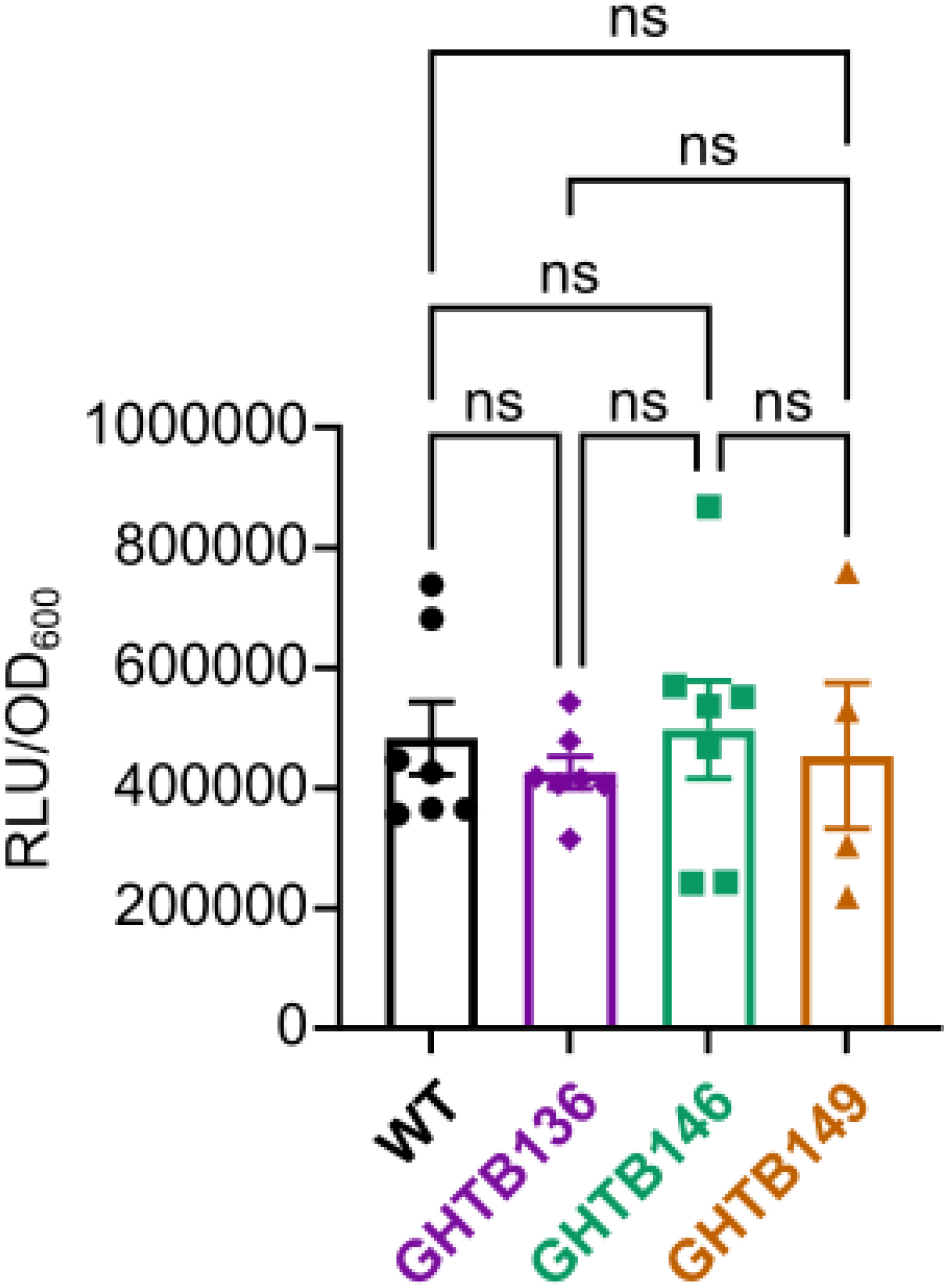
Basal ATP levels in C10-resistant mutants. The indicated strain of *Mtb* was cultured in Sauton’s liquid medium and ATP levels were measured by the BacTiter Glo assay. Data was collected from either 2 (GHTB149) or 3 (WT, GHTB136, GHTB146) independent experiments performed on separate days, n=4-7. The relative luminescence units (RLU) were normalized to the OD_600_ of the culture to control for differences in cell density. A one-way ANOVA was performed to determine if any of the strains had significantly different levels of ATP at baseline. ns, not significant.

**Figure S3:**
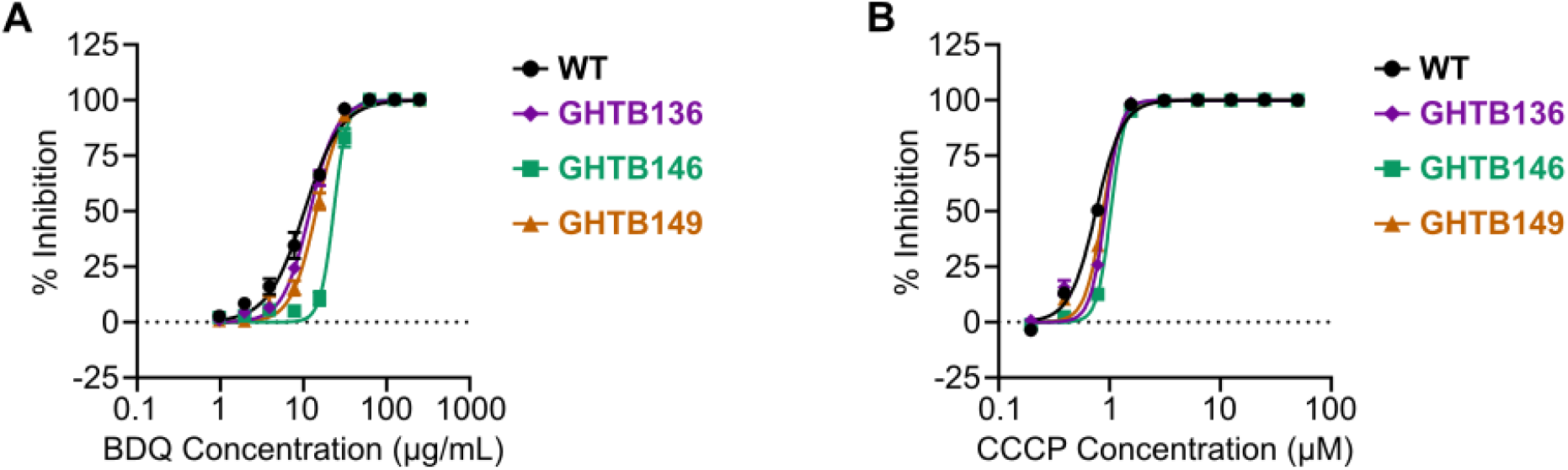
C10-resistant mutants are not cross-resistant to direct ETC inhibitors. (A-C) The indicated strain of *Mtb* was cultured in the presence of increasing concentrations of (A) BDQ or (B) CCCP for 1 week, and the % inhibition of *Mtb* growth and metabolism was determined using the resazurin assay, n=3. A best fit curve was determined in GraphPad Prism.

**Figure S4:**
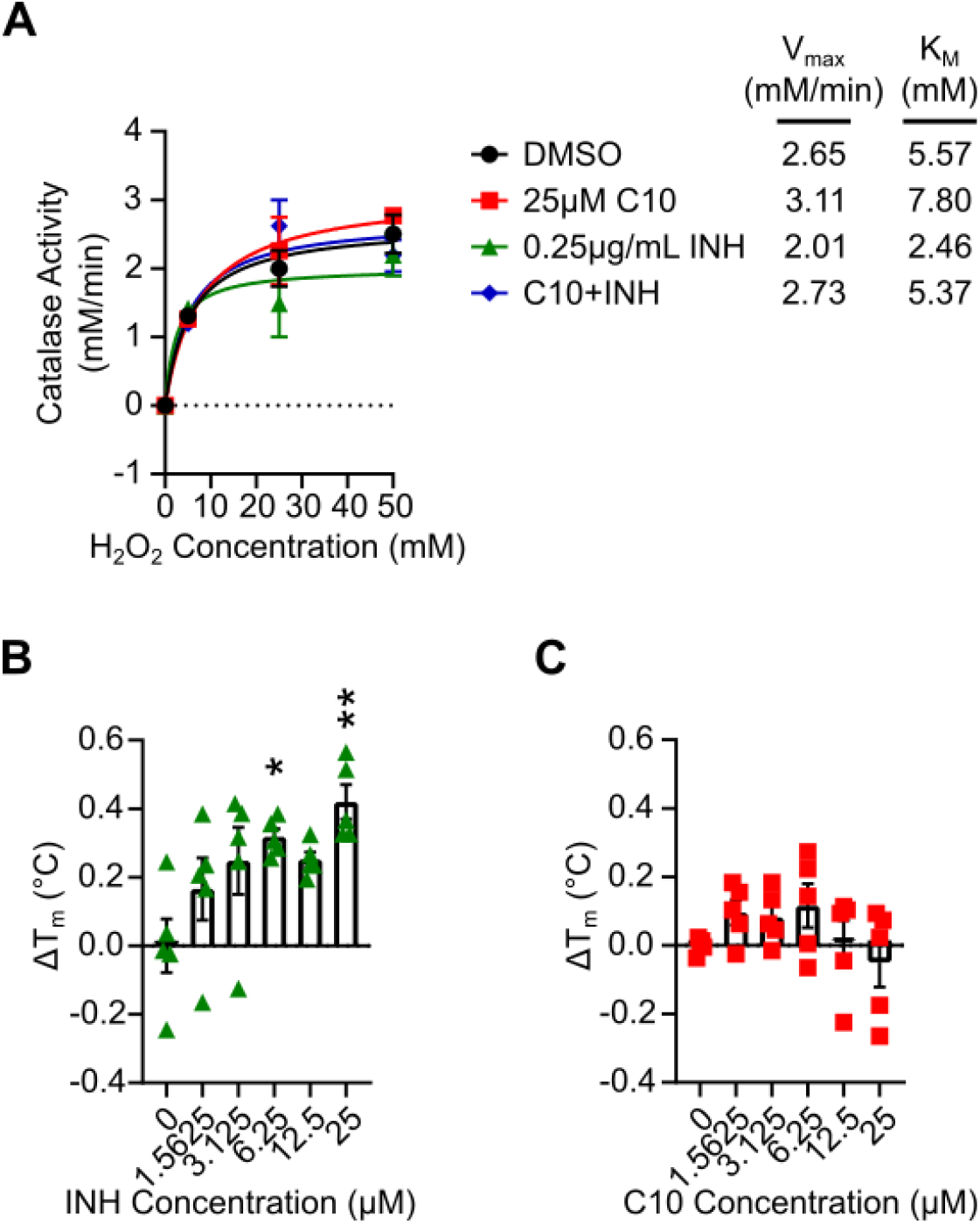
C10 does not increase KatG catalase activity or thermal stability. (A) Purified *Mtb* KatG protein (25nM) was incubated with or without 25μM C10 and/or 0.25μg/mL INH in 50mM potassium phosphate buffer pH 7.0, and the catalase activity, or H_2_O_2_ degradation, was monitored through a decrease in absorbance over time at λ_abs_=240nm in a UV/Vis spectrophotometer, n=3. The data in the graph was fitted with the Michaelis-Menten equation in GraphPad Prism to determine the V_max_ and K_M_ of KatG in each condition, which are indicated in the figure legend. (B-C) Purified *Mtb* KatG protein (1.1μM) was incubated with (B) INH or (C) C10 in 50mM potassium phosphate buffer pH 7.0 with 50μM MnCl_2_ before mixing samples with Sypro Orange, incubating at increasing temperatures, and monitoring fluorescence over time. The melting temperature (T_m_) was calculated by fitting curves with a Boltzmann sigmoidal equation in GraphPad Prism, and the ΔT_m_ was calculated as the difference between the T_m_ of each sample and the average of the untreated control samples (n=5). Statistically significant differences were identified by performing a 1-way ANOVA with Tukey’s post test, and data points that are statistically significantly different from the untreated control are indicated with stars. * P<0.05, ** P<0.01. For all pairwise comparisons see Supplementary Table S8.

**Figure S5:**
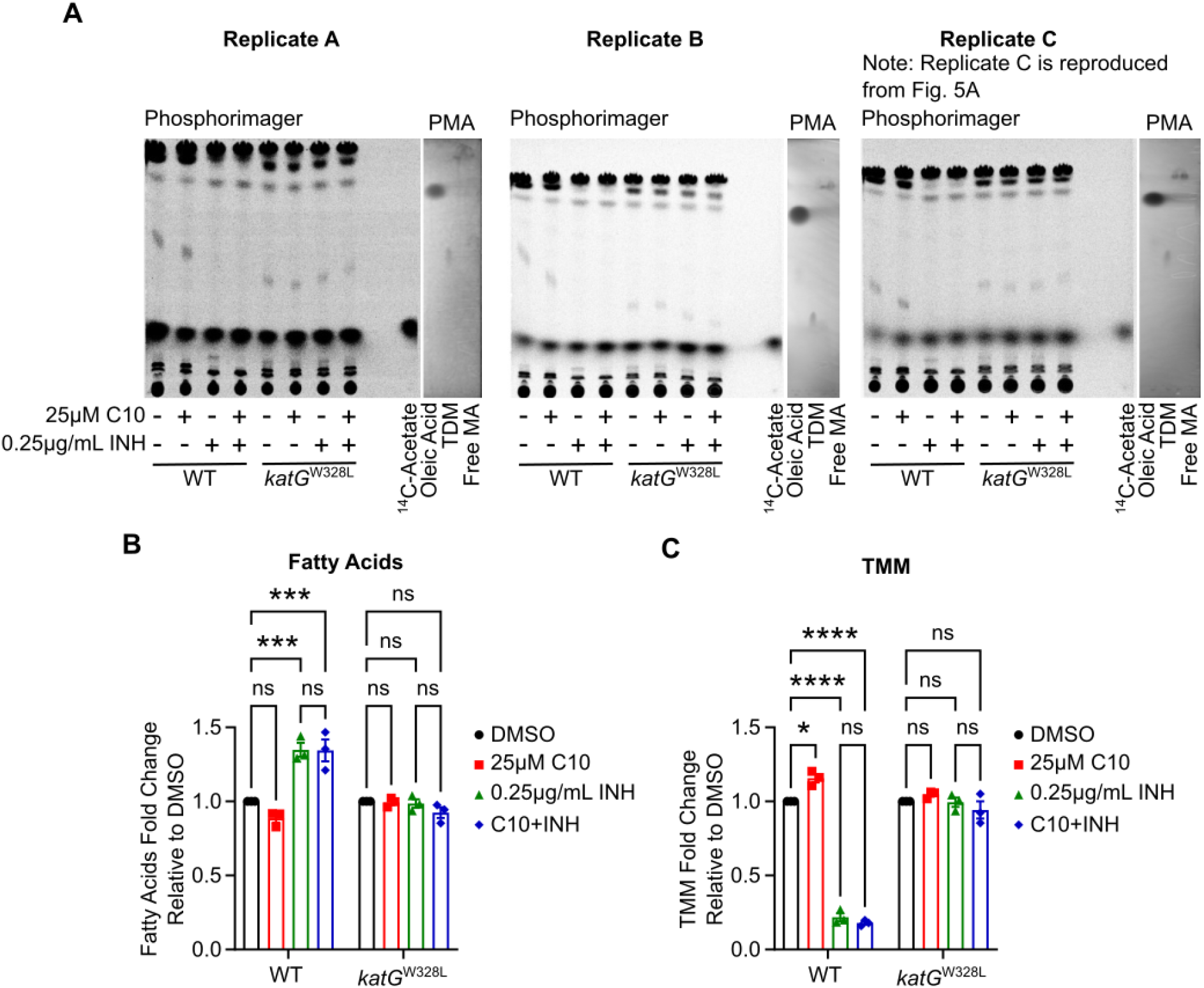
C10 does not alter mycolic acid biosynthesis in the presence or absence of INH. (A) WT or *katG*^W328L^ *Mtb* was cultured in Sauton liquid medium, treated with 25μM C10 and/or 0.25μg/mL INH, and immediately exposed to 2μCi/mL of ^14^C-labeled acetate. After 20 hours, lipids were extracted and analyzed by TLC to measure the *de novo* synthesis of mycolic acids and other lipids. The TLC plate was developed with 75:10:1 Chloroform:Methanol:H_2_O and radioactivity was analyzed by phosphorimaging. Bands corresponding to free mycolic acid (MA), fatty acid, trehalose dimycolate (TDM), and ^14^C-acetate were identified by comigration with a standard. Cold standards were imaged by coating the TLC plate in phosphomolybdic acid and charring. Trehalose monomycolate (TMM) is indicated with a * to emphasize that this lipid is putatively identified, and not correlated with a standard. (B-C) The intensity of bands corresponding to (B) TMM and (C) fatty acids were quantified in Image J, and normalized to the DMSO sample, with each WT sample being normalized to WT DMSO and each *katG*^W328L^ being normalized to *katG*^W328L^ DMSO in order to compare across replicates, n=3. Note that one of the images in panel A, is the same image that is presented in Figure 6 in the main text, reproduced here for comparison to the other replicates. Statistically significant differences were determined by 2-way ANOVA and select pairwise comparisons are depicted in the figure. ns not significant, *** P<0.001, **** P<0.0001. For all pairwise comparisons see Supplementary Table S9.

